# Cohesin organizes 3D DNA contacts surrounding active enhancers in *C. elegans*

**DOI:** 10.1101/2023.09.18.558239

**Authors:** Jun Kim, Haoyu Wang, Sevinç Ercan

**Affiliations:** Department of Biology, Center for Genomics and Systems Biology, New York University, New York, NY, USA

## Abstract

In mammals, cohesin and CTCF organize the 3D genome into topologically associated domains (TADs) to regulate communication between *cis*-regulatory elements. Many organisms, including *S. cerevisiae*, *C. elegans*, and *A. thaliana* contain cohesin but lack CTCF. Here, we used *C. elegans* to investigate the function of cohesin in 3D genome organization in the absence of CTCF. Using Hi-C data, we observe cohesin-dependent features called “fountains”, which are also reported in zebrafish and mice. These are population average reflections of DNA loops originating from distinct genomic regions and are ∼20-40 kb in *C. elegans*. Hi-C analysis upon cohesin and WAPL depletion support the idea that cohesin is preferentially loaded at NIPBL occupied sites and loop extrudes in an effectively two-sided manner. ChIP-seq analyses show that cohesin translocation along the fountain trajectory depends on a fully intact complex and is extended upon WAPL-1 depletion. Hi-C contact patterns at individual fountains suggest that cohesin processivity is unequal on each side, possibly due to collision with cohesin loaded from surrounding sites. The putative cohesin loading sites are closest to active enhancers and fountain strength is associated with transcription. Compared to mammals, average processivity of *C. elegans* cohesin is ∼10-fold shorter and NIPBL binding does not depend on cohesin. We propose that preferential loading and loop extrusion by cohesin is an evolutionarily conserved mechanism that regulates the 3D interactions of enhancers in animal genomes.

## Introduction

In mammalian genomes, cohesin and CTCF form topologically associating domains (TADs) (da Costa-Nunes & Noordermeer, 2023). The earlier models of TAD formation encompassed three main features, cohesin is uniformly loaded, cohesin performs two-sided loop extrusion, and loop extrusion is stalled at CTCF-binding sites located at TAD boundaries (Fudenberg et al., 2017). The second and the third assumptions of the model were visualized *in vitro* using single-molecule assays, thus bridging the gap between the theoretical model and the experimental observation *in vivo* (Davidson et al., 2019; Kim et al., 2019). Further studies refined the earlier models and proposed mechanistic variations of two-sided loop extrusion (e.g. frequent switching of direction in a one-sided extrusion scheme), effect of cohesin molecules meeting one another (resulting in bypassing to form secondary folds in loops or colliding and stopping extrusion on one side), and specific proteins controlling loop extrusion as loader/facilitator (NIPBL), remover (WAPL), or acting as barriers (e.g. MCM complex, RNA Pol II, DNA topology) (Banigan et al., 2023; Corsi et al., 2023; Dequeker et al., 2022; Morao et al., 2022).

Recent studies also required an update to uniform loading assumption, because Hi-C detected features termed “jets”, “plumes” or “fountains” depicted as secondary diagonals protruding from narrow segments suggest cohesin to be loaded at specific sites (Guo et al., 2022; Shao et al., 2024; Wike et al., 2021). The preferential loading sites locally increase the frequency of cohesin-mediated loops, thus providing another layer of 3D genome folding with potential to regulate contacts between *cis*-regulatory elements. Such a mechanism of genome folding could have strong regulatory implications for species that contain tissue specific enhancers but lacking CTCF, such as *Caenorhabditis elegans*.

*C. elegans* is a small nematode with ∼100 Mb genome and ∼20,000 genes (Heger et al., 2009). Median gene length is ∼2 kb and intergenic distances between genes (excluding operons encompassing ∼15% of genes) range from ∼2 to 10 kb (Allen et al., 2011; Girard et al., 2007; Nelson et al., 2004). Like other species, enhancers are distinguished from promoters based on enrichment of distinct and conserved histone modifications, including H3K27 acetylation and H3K4 methylation (Daugherty et al., 2017; Evans et al., 2016). Mapping enhancer chromatin features and analyses of individual genes suggest that most enhancers are located ∼ 2 kb from promoters, but some enhancers can be up to ∼12 kb away (Shi et al., 2009).

In humans, somatic cohesin is composed of four subunits, SMC1, SMC3, RAD21, and STAG (Hassler et al., 2018). The STAG subunit has two isoforms STAG1 and STAG2 that forms cohesin in mutually exclusive manner (Alonso-Gil et al., 2023). Several subunits of mammalian cohesin are specific for meiosis (Ishiguro, 2019). In *C. elegans*, the three of the four subunits of cohesin, SMC-1, SMC-3, and SCC-3 (STAG), has a single copy, while the kleisin subunit has 5 variants: COH-1, COH-2 (aka SCC-1), COH-3, COH-4, and REC-8 (Severson & Meyer, 2014; Wood et al., 2010). COH-2, but not COH-1, has mitotic functions, while COH-3/4 (paralogs) and REC-8 have meiotic functions (Hernandez et al., 2018; Mito et al., 2003; Severson & Meyer, 2014; Yu et al., 2023). We reasoned that analyses of cohesin binding and 3D genome organization upon rapid depletion of SMC-3 and its negative regulator WAPL-1 would reveal cohesin function in *C. elegans* genome organization.

In our study of *C. elegans* 3D genome, we observe and identify “fountains” on the Hi-C matrix, which are reflections of increasing lengths of DNA loops up to ∼20-40 kb in length originating from distinct sites. The fountains are lost and extended upon acute (1 hour) depletion of SMC-3 and WAPL, respectively. In agreement with Hi-C data, ChIP-seq signals of cohesin subunits SMC-3 and SMC-1 follow the trajectory of the fountains, extending upon WAPL depletion. In contrast, the binding patterns of the NIPBL, positive regulator of cohesin, remained unchanged upon SMC-3 or WAPL depletion, suggesting that NIPBL does not move with cohesin in *C. elegans*. Fountains are associated with highly transcribed regions and the origins are closest to active enhancers. Despite the elimination of 3D DNA contacts forming the fountains upon acute depletion of cohesin, a similar degree of change in transcription was not observed in the RNA Pol II ChIP-seq or mRNA-seq data. Together, our results are consistent with a model that NIPBL is recruited at or near active enhancers, resulting in increased loading of cohesins. The preferentially loaded cohesin translocates along the DNA via two-sided loop extrusion. The transcriptional program is robust to acute perturbations to 3D DNA contacts surrounding the enhancers.

## Materials and Methods

### Worm strain and growth

Worms were grown and maintained at 20-22°C on Nematode Growth Medium (NGM) plates containing *E. coli* strains OP50-1 and/or HB101. To isolate synchronized L2/L3 worms, gravid adults were bleached in 0.5 M NaOH and 1.2% bleach, and embryos were hatched overnight in M9 buffer (22 mM KH2PO4 monobasic, 42.3 mM Na2HPO4, 85.6 mM NaCl, 1mM MgSO4). The resulting starved L1s were grown for 24 hours at 22°C. Degron-GFP tagged alleles were produced by SUNY Biotech and crossed to CA1200 strain expressing TIR1 under the control of eft-3 promoter to produce ERC102 syb5520 [smc-3::GGGGS::AID::emGFP] III, [eft-3p::TIR1::mRuby::unc-54 3’UTR + Cbr-unc-119(+)] II, ERC103 syb6035 [wapl-1::GGGGS::AID::emGFP] IV, [eft-3p::TIR1::mRuby::unc-54 3’UTR + Cbr-unc-119(+)] II and ERC107 syb8826[scc-2::GGGGS::AID], ieSi57 II; unc-119(ed3) III.

### Auxin treatment

Auxin (indole-3-acetic-acid, Fisher 87-51-4) was resuspended in 100% ethanol to a concentration of 400 mM. Plates were prepared by adding resuspended auxin at a concentration of 1 mM to NGM media before pouring. Synchronized L2/L3 worms were washed three times with M9 and split into two. Half of the worms were transferred to NGM 10 cm plates containing 1mM of auxin. The other half were placed in normal NGM 10 cm plates (no-auxin control). A maximum of 300 μL of settled worms was placed in one 10 cm plate. Worms were then washed one time with M9 and processed according to future application. For ChIP and Hi-C, worms were crosslinked in 2% formaldehyde for 30 minutes, followed by quenching in 125mM glycine for 5 minutes, one wash with M9 and two washes with PBS, PMSF and protease inhibitors. For RNA-seq, worms were stored in Trizol.

### Microscopy

*The* worms were washed off the plate in M9 buffer, crosslinked in 0.5% formaldehyde for 15 minutes and ‘freeze cracked’ by submerging in liquid nitrogen for 1 minute and then thawing in a 37°C water bath. The worms were washed twice using 70% ethanol followed by a third wash using the wash buffer (PBS-0.05% Tween-20). The worms were resuspended in 1mL of wash buffer and 1ul of 2ug/mL DAPI (FisherEN62248) and were incubated in 37°C water bath for 30 minutes. The worms were then washed twice in PBS and imaged on slides for DAPI and GFP.

### ChIP-seq

Two biological replicates with matching input samples were performed for each experiment. Around 100 μL of L2/L3 larvae were dounce-homogenized with 30 strokes in FA buffer (50 mM HEPES/KOH pH 7.5, 1 mM EDTA, 1% Triton X-100, 0.1% sodium deoxycholate, 150 mM NaCl) supplemented with PMSF and protease inhibitors (Calbiochem, 539131). Dounced worms were sonicated in 0.1% sarkosyl for 15 minutes using a Picoruptor to obtain chromatin fragments between 200 and 800 bp. Protein concentration was determined using Bradford assay (Biorad 500-0006). 1-2 mg of protein extract was used per ChIP and 5% was taken out to use as input DNA. The remaining protein extract was incubated with 3 to 10 ug of antibody at 4°C rotating overnight in a volume of 440μL. 40μL of (1:1V) Protein A and/or G Sepharose beads that were previously washed 3 times with FA buffer, were added and incubated, rotating at 4°C for 2 hours. Beads were washed with 1mL of each of the following buffers: 2 times with FA buffer, 1 time with FA-1mM NaCl buffer, 1 time with FA-500mM NaCl buffer, 1 time with TEL buffer (0.25 M LiCl, 1% NP-40, 1% sodium deoxycholate, 1 mM EDTA, 10 mM Tris-HCl, pH 8.0) and 2 times with TE buffer. Immunoprecipitated chromatin was eluted from beads by incubating in ChIP elution buffer (1% SDS, 250 mM NaCl, 10 mM Tris pH 8.0, 1 mM EDTA) at 65°C for 30 minutes, treated with Proteinase K and reverse crosslinked at 65°C overnight. Half of the ChIP DNA and 30 ng of input DNA were used for library preparation. End repair was performed in T4 ligase reaction buffer (New England Biolabs, NEB), 0.4mM of dNTPs, 20 U of T4 Polynucleotide kinase (NEB), 3.5 U of Large (Klenow) fragment (NEB) and 6 U of T4 DNA polymerase for 30 minutes at 20°C in a total volume of 33 μL. Reaction was cleaned using Qiagen MinElute PCR purification kit. A-tailing reaction was performed in NEB buffer 2, 0.2 mM of dATPs and 7.5 U of Klenow fragment-exo (NEB) at 37°C for 60 minutes in 25 μL. Reaction was cleaned using Qiagen MinElute PCR purification kit. Illumina TruSeq adapters were ligated to DNA fragments in a reaction containing 2X Quick ligation buffer (NEB), 0.25 μM of adapter and 2 μL of Quick ligase (NEB) for 20 minutes at 23°C in 40 μL. Reaction was cleaned using Agencourt AMPure XP beads and the eluted DNA was PCR amplified in 50 μL using Phusion Polymerase and TruSeq primers, forward: AAT GAT ACG GCG ACC ACC GAG ATC TAC ACT CTT TCC CTA CAC G∗A, reverse: CAA GCA GAA GAC GGC ATA CGA GA∗T, where ∗ represents a phosphorothioate bond. PCR reactions were cleaned using Qiagen MinElute PCR purification kit. The eluted DNA was run on a 1.5% agarose gel and fragments between 250-600 bp were gel extracted using Qiagen gel extraction kit. Library concentration was determined using KAPA Library quantification kit. Single-end 75 bp sequencing was performed using the Illumina NextSeq 500.

ChIP-seq data analysis: Bowtie2 version 2.4.2 was used to align 75 bp single-end reads to WS220 with default parameters (Langmead & Salzberg, 2012). Bam sorting and indexing was performed using samtools version 1.11 (Danecek et al., 2021). BamCompare tool in Deeptools version 3.5.0 was used to normalize for the sequencing depth using CPM and create ChIP-Input coverage with a bin size of 10 bp and 200 bp read extension (Ramirez-Gonzalez et al., 2012). Only reads with a minimum mapping quality of 20 were used, and mitochondrial DNA, PCR duplicates, and blacklisted regions were removed (Amemiya et al., 2019). The average coverage data was generated by averaging ChIP-Input enrichment scores per 10 bp bins across the genome. ChIP-seq peaks are identified using MACS2 version 2.1.1 (https://github.com/macs3-project/MACS). Peaks are called using both the individual replicates and combined bam files with minimum false discovery rate of 0.05. The peaks shown below average ChIP-seq tracks are peaks called using the combined bam files, whereas the peaks shown below individual replicates are peaks called using the corresponding replicate.

### Hi-C

Two biological replicates were performed for each experiment. Crosslinked L2/L3 worms collected as described above were resuspended in 20 ul of PBS per 50ul of worm pellet, then dripped into liquid nitrogen containing mortar. The worms were grounded with pestle until fine powder. Grounded worms were crosslinked again in 2% formaldehyde using the TC buffer as described by the Arima High Coverage Hi-C kit, which uses four 4-base cutters, DpnII, HinfI, DdeI, and MseI. The Arima manufacturer’s protocol was followed including the recommended method of the library preparation using KAPA Hyper Prep Kit. Paired-end 100 bp sequencing was performed using the Illumina Novaseq 6000.

Hi-C data analysis: The Hi-C data was mapped to ce10 (WS220) reference genome using default parameters of the Juicer pipeline version 1.5.7 (Durand et al., 2016). The biological replicates were combined using juicer’s mega.sh script. The mapping statistics from the inter_30.txt output file are provided. The inter_30.hic outputs were converted to cool format using the hicConvertFormat of HiCExplorer version 3.6 (Ramirez et al., 2018; Wolff et al., 2018) in two steps using the following parameters: 1) --inputFormat hic, --outputFormat cool, 2) -- inputFormat cool --outputFormat cool --load_raw_values. The cool file was balanced using cooler version 0.8.11 using the parameters: --max-iters 500, --mad-max 5, --ignore-diags 2 (Abdennur & Mirny, 2020). The balanced cool file was used for all downstream analysis.

For computing log-binned P(s) and its log-derivative cooltools version 0.4.0 (https://github.com/open2c/cooltools) was used. For visualizing ChIP-seq data with Hi-C data in python, pyBigwig version 0.3.18 (https://github.com/deeptools/pyBigWig) was used.

Fountain identification: Fountains must be (1) observed in control, (2) observed in WAPL depletion conditions, and (3) show significant decrease upon cohesin depletion. We apply the tool implemented in cooltools (version 0.4.0), which was originally developed to identify elements whose interaction across is insulated, to instead identify elements whose interaction across is facilitated. Instead of using local minima, the local maxima were used. Peak prominence was calculated for local maxima to identify their strength relative to the surrounding. Then otsu’s threshold was used to remove local maxima that arises from random noise. We identified 449 (criteria - 1) and 587 (criteria - 2) local maxima in control condition and WAPL depletion condition. To identify elements whose interaction across is significantly lost upon cohesin depletion, we used insulation score delta (control minus SMC3 depletion) to identify local maxima. The identified 831 (criteria - 3) significant local maxima were further subsetted based on their overlap with criteria 1 and 2, resulting in the final 287 fountains (Supplemental Fig S1C, D). The ‘facilitation strength’ is the peak prominence value of insulation delta (control minus SMC3 depletion). We reason that this metric best captures how strongly the interaction across the element is facilitated by cohesin.

Fontanka (https://github.com/agalitsyna/fontanka) (Aleksandra Galitsyna et al., 2023) was used with the following steps. (i) Snippets extraction: we manually identified three different types of shapes based on the tilt of the fountain: center, left, and right. We manually selected 10 snippets that appeared to show left or right tilt. For each shape, the 10 snippets were used to generate observed-over-expected on-diagonal pile-up matrices with flanking +/-100kb window. This pile-up is used as the ‘reference mask’ to which the snippets from the rest of the genome is compared to. (ii) Fountain score and Scharr score: we used the fontanka slice-windows function to generate genome-wide snippets using the parameter -W 100_000 for both control condition and WAPL depletion conditions using 2 kb binned matrix. Fountain and Scharr score for each window were calculated by Fontanka-apply-fountain-mask. (iii) Peak calling and Thresholding: we applied otsu thresholding method on peak prominence returned from fontanka-apply-fountain-mask function and applied a minimum threshold of 0.2 for fountain score. Scharr score, which measures the uniformity/sharpness of the Hi-C matrix, can be used to infer rearrangements and incomplete reference genome data. Windows of Hi-C matrix with sharp transitions, which often arises from imperfect reference genome (rearrangements and missing segments), have higher Scharr score, whereas windows containing smooth transitions in contact scores have lower Scharr score. We only removed the top scoring 5% of candidate fountains (false positives due to reference genome) based on previous work showing that long-read assembly of *C. elegans* genome only added 1.8 Mb (<2%) to the existing reference genome (Yoshimura et al., 2019). (iv) Leveraging WAPL condition: we kept control fountains that overlapped (+/- 10 kb) with those called in the WAPL depletion which shows sharper contrast in fountain shapes. (vi) Validation: we ensured that the average on-diagonal pile-up of the called fountains shows distinct shapes.

### mRNA-seq

L3 larvae were collected and stored in Trizol (Invitrogen) at -70°C. Total RNA was extracted through the freeze-crack method and processed as described previously (Albritton et al., 2014). RNA was cleaned up using Qiagen RNeasy MinElute Cleanup kit. mRNAs were purified using Sera-Mag Oligo (dT) beads (Thermo Scientific), then fragmented using Ambion (AM8740), and cleaned using Qiagen RNAeasy minElute kit. RNA was reverse transcribed using Superscript III system (Invitrogen, 18080-051). Residual dNTP was removed using ethanol precipitation. The second strand was synthesized using DNA polymerase I (Invitrogen). dsDNA was purified using Qiagen DNA minElute kit. Single-stranded illumine libraries were prepared as indicated for ChIP-seq with the following modification: prior to PCR amplification, uridine was digested using Uracil-N-Glycosylase (ThermoScientific EN0361).

mRNA-seq data analysis: Single-end 75 bp sequencing was performed using the Illumina NextSeq 500. Reads were mapped to WS220 (ce10) *C. elegans* genome using HISAT2 version 2.2.1 (Kim et al., 2015) with the parameter --rna-strandness R. The SAM output from HISAT2 and WS220 gtf file were used to calculate counts per gene using HTSeq version 0.13.5 (Anders et al., 2015). The raw counts were normalized FPKM using cufflinks version 2.2.1 (Roberts et al., 2011), and then FPKM was converted to TPM. For comparing between conditions, the raw counts were used for the R package DESeq2 version 1.30.0 (Love et al., 2014).

### Sequencing data and analysis script availability

The data are available at the Gene Expression Omnibus under GSE237663. A list of data and statistics are provided in Supplemental File 1. Analysis scripts are provided at https://github.com/ercanlab/2023_Kim_cohesin.

## Results

### Fountains are a feature of 3D genome organization detected by Hi-C

*C. elegans* genome is organized into six similar sized chromosomes, including five autosomes (I-V) and one X-chromosome. The chromosomes are holocentric and are partitioned into three. The gene-rich central regions make inter-chromosomal contacts, and gene-poor left and right arms are frequently located close to the nuclear lamina (Bian et al., 2020; Ho et al., 2014; Liu et al., 2011). At the mega-base scale, the autosomes display A/B compartmentalization without loop-anchored TADs, which is consistent with the lack of CTCF (Figure 1). The X-chromosomes contain weaker compartments along with loop-anchored TADs, formed by a condensin I variant that functions in dosage compensation (DC) in hermaphrodites (Anderson et al., 2019; Bian et al., 2020; Crane et al., 2015; Kim et al., 2022; Morao et al., 2022; Sawh & Mango, 2022). At the multi-kilobase scale, we observe contact enrichment patterns that orthogonally protrude from the main-diagonal across all chromosomes (Figure 1).

**Figure 1.**
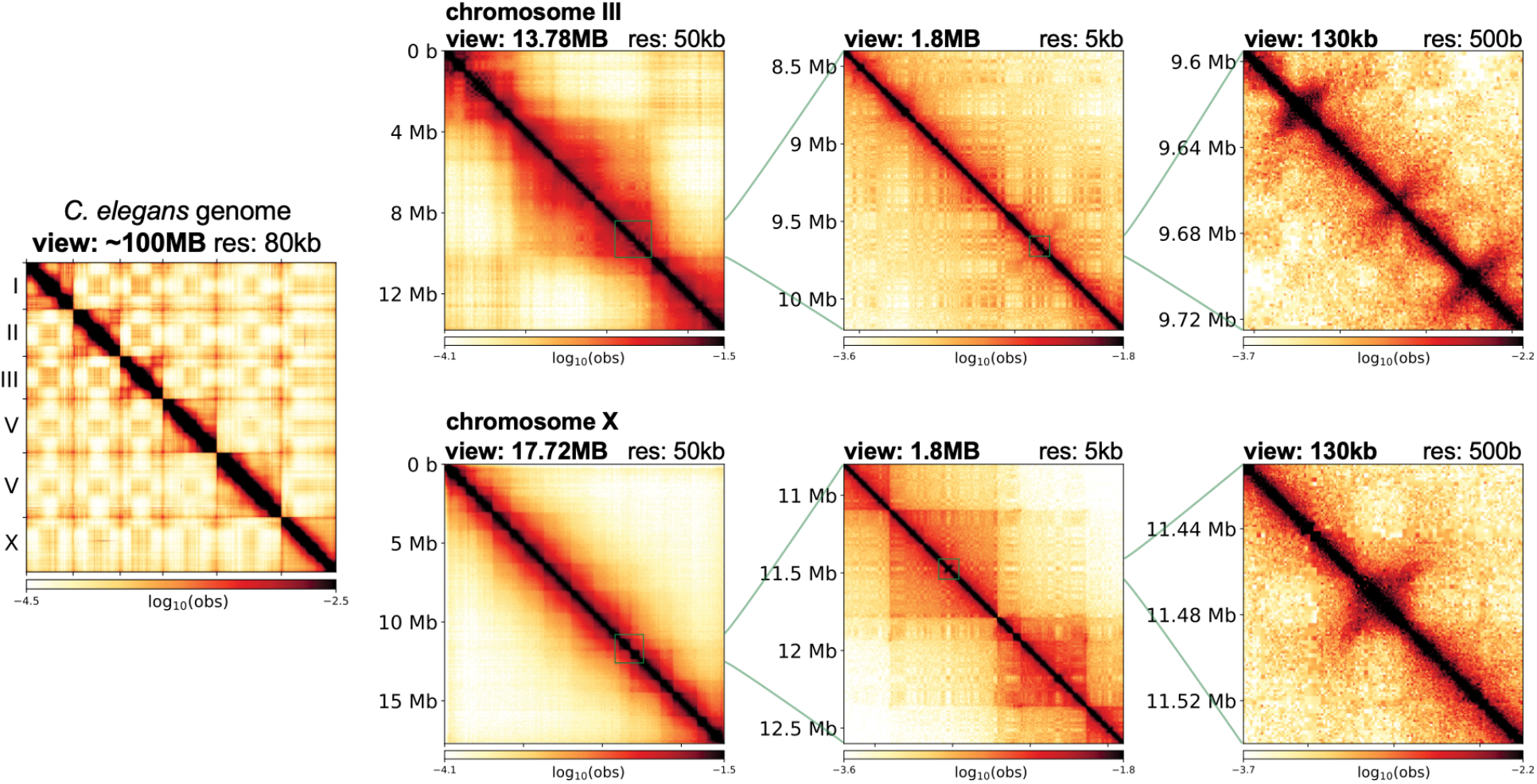
3D organization of the *C. elegans* genome at different scales. Leftmost panel shows the Hi-C contact matrix of the entire genome. The right three columns show an example of an autosome (top row) and X-chromosome (bottom row) at different scales. At chromosome-wide scale (first column), autosomes show clear separation between the two interacting flanking arms and the center, whereas the X-chromosome is more uniform. At the mega-base scale (second column), autosomes show checkerboard pattern, whereas X-chromosome additionally harbors TADs. At kilo-base scale (last column), both autosomes and X-chromosome show fountains that appear as protruding 3D DNA contacts from the main diagonal.

DNA loops originating and extending symmetrically in both directions from a specific site in a population of cells form enriched Hi-C contacts perpendicular to the main-diagonal. Such patterns were termed “chromatin jets” in mice (Guo et al., 2022). Recent studies expanded on similar Hi-C features and defined ‘fountains,’ which are not perfectly perpendicular and show “fanning”, due to heterogeneity of the formed loops (A. Galitsyna et al., 2023; Isiaka et al., 2023; Shao et al., 2024). In the absence of quantitative criteria that distinguish chromatin jets and fountains, in our *C. elegans* Hi-C data we will refer to the contacts protruding from the main-diagonal as “fountains” to be more inclusive of fanning.

To test whether the fountains are due to loop extrusion by cohesin, we decided to perform Hi-C analysis in the absence of cohesin or WAPL. While cohesin depletion is sufficient to test for requirement, we reason that WAPL depletion is necessary to distinguish loop extrusion from other mechanisms. For instance, if the fountains were formed by cohesin without gradual increase in the DNA loop size, increasing residence time of cohesin would result in more pronounced fountains of the same size. In contrast, the loop-extrusion model predicts that WAPL depletion leads to cohesin translocating farther, extending the DNA loops, thus fountains in length.

### Acute depletion of SMC-3 and WAPL from somatic cells

In *C. elegans*, there is a singular subunit for SMC-1, SMC-3, and SCC-3 (Figure 2A). To study the effect of cohesin loss, we targeted SMC-3 subunit for auxin inducible degradation. WAPL is encoded by *wapl-1* in *C. elegans* (Gandhi et al., 2006; Kueng et al., 2006). The two proteins, SMC-3 and WAPL were C-terminally tagged with degron-GFP at the endogenous locus for auxin inducible depletion, as employed previously (Morao et al., 2022; Zhang et al., 2015). To enrich post-mitotic interphase cells, we used the early L3 stage, when most somatic cells have stopped dividing and the germline has not significantly proliferated (Sulston & Horvitz, 1977). Furthermore, we depleted the proteins only in somatic cells, using a strain in which the auxin response gene TIR1 is under the control of a soma-specific promoter *eft-3* (Zhang et al., 2015). This background strain is used as a control where TIR1 is expressed but none of the endogenous genes are tagged with degron-GFP (Figure 2B).

**Figure 2.**
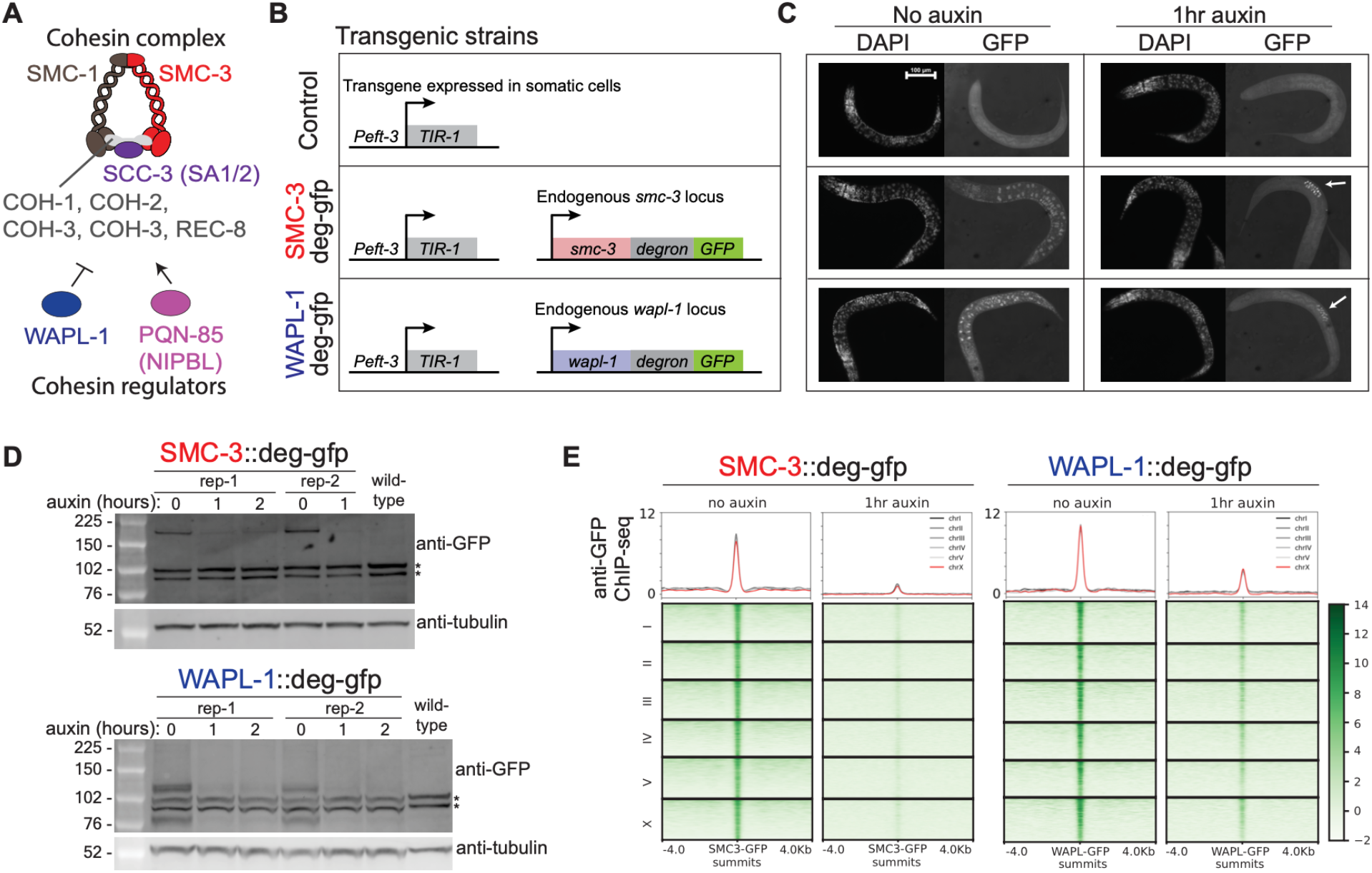
Acute depletion of cohesin and WAPL from somatic cells. A) *C. elegans* subunits of cohesin and its regulators. B) Transgenic strains used in this paper. Exogenously inserted TIR1 is expressed in somatic cells using the *eft-3* promoter. Endogenous *smc-3* or *wapl-1* are C-terminally tagged with degron-gfp. C) DAPI and GFP images of L3 stage worms. White arrow indicates that the proteins in the germline are not degraded upon auxin treatment due to TIR1 presence only in the somatic cells. Scale bar: 100um. D) Western blot images of L3 stage worms with increasing auxin treatment duration. Anti-GFP antibody was used to show global depletion of the tagged protein. Anti-tubulin antibody was used as loading control. Asterisk indicates background bands. E) Meta-plot of ChIP-seq data grouped into different chromosomes. Anti-GFP antibody was used to show uniform depletion of SMC-3 and WAPL.

Upon visualizing DAPI and GFP, we observe that in the control condition, both cohesin and WAPL show ubiquitous GFP signals, suggesting that cohesin and WAPL are present in the interphase as well as the few germ cells present in early L3 larvae (Figure 2C, no auxin). Upon auxin treatment, cohesin and WAPL are noticeably depleted in somatic cells but are maintained in the germ cells as expected from somatic expression of TIR1 (Figure 2C, 1hr auxin). At population average, western blot analysis of degron-GFP tagged proteins show noticeable depletion in the total amount of protein (Figure 2D). This depletion is maximally achieved by 1-hour treatment, and no further depletion was observed upon 2-hour treatment. Therefore, we chose 1-hour as the ‘depletion condition.’ Lastly, we performed ChIP-seq using anti-GFP and observed depletion across the genome (Figure 2E). In summary, both SMC-3 and WAPL are acutely depleted from somatic cells with 1-hour auxin treatment.

### DNA loops forming the fountains are cohesin dependent and extend upon WAPL depletion

Fountains are heterogeneous. They can be left-tilted (Figure 3A), right-tilted (Figure 3B) or show complex trajectories (Supplemental Fig S1A). In all cases, SMC-3 depletion eliminated and WAPL depletion extended fountains. For example, in the control condition, we observe a short fountain (Figure 3A, left). In the SMC-3 depletion, the fountain is lost, indicating that it is cohesin dependent (Figure 3A, center). In the WAPL depletion, the trajectory of the fountain is extended, suggesting that cohesin initially forms a smaller loop and then a larger loop with increasing residence time on DNA (Figure 3A, right). The tilt of the fountains indicates that the processivity of cohesin is not equal on each side (Figure 3C).

**Figure 3.**
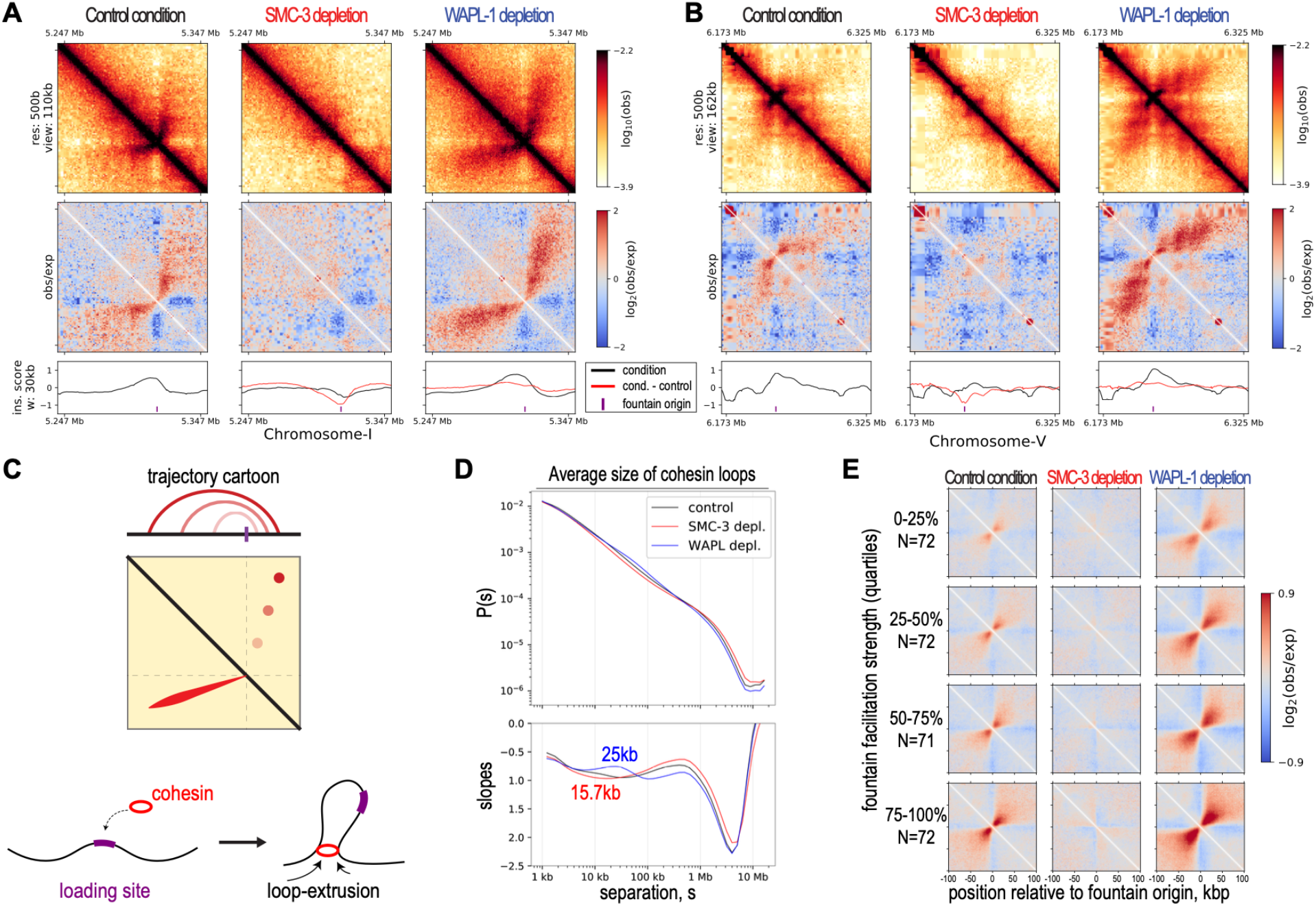
Cohesin forms fountains via loop-extrusion. A) Example Hi-C snapshot of left-directed fountain in control, SMC-3, and WAPL depletion conditions. Going from the top to bottom row: The observed balanced matrix, distance-decay normalized observed-over-expected matrix, and insulation score using 30kb-window and 500bp-step (black lines – condition (control of depletions), red lines – condition minus the control. The purple vertical line indicates identified fountain origins. B) Example Hi-C snapshot of right-directed fountain. C) Cartoon trajectory of left-directed fountain (panel A) arising from two-sided loop extruding factor reeling in DNA faster on the left than right side. D) Contact probability, P(s), as a function of linear genomic separation between pairs of loci, s (top row), and its log-derivative, the slopes of P(s) in log-space, (bottom row) are plotted for three conditions in different colors. The inferred average loop size for cohesin in unperturbed state (red) and the extended average loop size for cohesin in WAPL depletion (blue) are annotated. E) On-diagonal pileup of observed-over-expected matrix centered at the fountain origins grouped into four quantiles based on the facilitation strength.

*In vitro*, cohesin performs two-sided loop extrusion (Davidson et al., 2019; Kim et al., 2019). In simulation, cohesin was inferred to be an *effectively* two-sided loop extruding factor (Banigan et al., 2020). Hi-C data cannot distinguish if cohesin acts as a single molecule pulling DNA at both ends or two tethered molecules each extruding from one end. Regardless, our observation that cohesin mediates DNA contacts protruding from the Hi-C diagonal supports the conclusion that cohesin effectively functions as a two-sided loop extruding factor *in vivo*. Furthermore, detection of cohesin loops in the Hi-C assay supports the idea that they are formed due to preferential cohesin loading at stereotypical locations.

### Cohesin-mediated DNA loops in C. elegans are shorter than those observed in mammals

While fountains are pronounced Hi-C features, they may reflect above-average cohesin behaviors on chromatin. The contact probability curve, P(s), and its log-derivative are powerful metrics for estimating the average loop size of the chromosome (Gassler et al., 2017; Polovnikov et al., 2023). In the mammals, this approach revealed that cohesin forms ∼100-200 kb loops on average (Davidson et al., 2019; Haarhuis et al., 2017; Polovnikov et al., 2023). Here, we use a similar metric to analyze the size of loops formed by cohesin. While we do not observe a local maximum in the log-derivative of P(s) that disappears in the SMC-3 depletion condition, we observe a local maximum that appears in WAPL depletion at ∼25 kb on autosomes (Figure 3D, Supplemental Fig S1B). We infer that the average loop size of cohesin in control condition is likely smaller and may correspond to local minima that becomes more pronounced in SMC-3 depletion, which is near ∼15 kb on autosomes. Interestingly, the cohesin mediated DNA loops are shorter on the X compared to autosomes (Supplemental Fig S1B). This may be related to the activity of a dosage compensation specific condensin forms TADs and represses transcription (Albritton & Ercan, 2018; Kim et al., 2022). We conclude that the loops formed by *C. elegans* cohesin are ∼10 fold smaller than the ones formed by the mammalian cohesin.

### Cohesin binding tracks the trajectory of the fountains and extends farther upon WAPL depletion

Our Hi-C analysis is consistent with a model of cohesin preferentially loading and loop extruding to form fountains. An orthogonal approach to test if the observed fountains are formed by the loop extruding cohesin is to analyze the distribution of cohesin with respect to the fountain origins. To identify fountains genome-wide, we determined sites across which DNA contacts are *facilitated by cohesin*. By our definition, cohesin-dependent fountains must be present in both control and WAPL depletion conditions and show significant decrease upon SMC-3 depletion (Supplemental Fig S1C). A total of 287 sites satisfied the criteria. Meta analysis of the average Hi-C signal in quartiles of “facilitation strength” (see methods) confirmed the identification of cohesin dependent fountain origins (Figure 3E) and was consistent between experimental replicates (Supplemental Fig S1D).

In simulation, loop extruding factors with a preferential loading site generates a ‘spreading’ pattern, where binding gradually decreases with increasing genomic distance from a loading source (Brandao et al., 2021). Loop extruders with known recruitment mechanisms support this pattern. ChIP-seq enrichment of bacterial condensin spreads out from the *parS* sequences (Sullivan et al., 2009; Wang et al., 2017; Wang et al., 2018). Similarly, *C. elegans* condensin DC shows reduced binding with increasing distance from the endogenous or ectopically inserted recruitment elements on the X (*rex* sites) (Kim et al., 2022; Morao et al., 2022). Targeted loading of cohesin using DNA damage or tetO-tetR system also show spreading of cohesin to nearby regions (Arnould et al., 2021; Han et al., 2023). Therefore, if cohesin is preferentially loaded at fountain origins, we predict cohesin binding to show a spreading pattern. Furthermore, under WAPL depletion, where the fountain length increases, the spreading pattern of cohesin would widen as cohesin travels farther away.

To analyze cohesin binding, we performed ChIP-seq for two subunits of the cohesin complex, SMC-1 and SMC-3. In the control condition, we observe that cohesin binds broadly for the span of approximately 20-kb region (Figure 4A, Supplemental Fig S2, black tracks). Upon SMC-3 depletion, both SMC-3 and SMC-1 are largely depleted (Figure 4A, red tracks). For SMC-1, we observe residual binding at the fountain origins (Figure 4B), which is consistent across biological replicates (Supplemental Fig S3A) and unlikely due to differences in the specificities of SMC-3 and SMC-1 antibodies to the fountain origins (Supplemental Fig S3B). The residual SMC-1 may reflect the process of searching through its potential target sites (Izeddin et al., 2014) or transient association as observed with the cohesin complex with defective ATPase activity (Ladurner et al., 2014). The loss of SMC-1 and SMC-3 in the flanking regions is consistent with the loss of the fountain in the Hi-C data (Figure 3), demonstrating that cohesin is essential for fountain formation.

**Figure 4.**
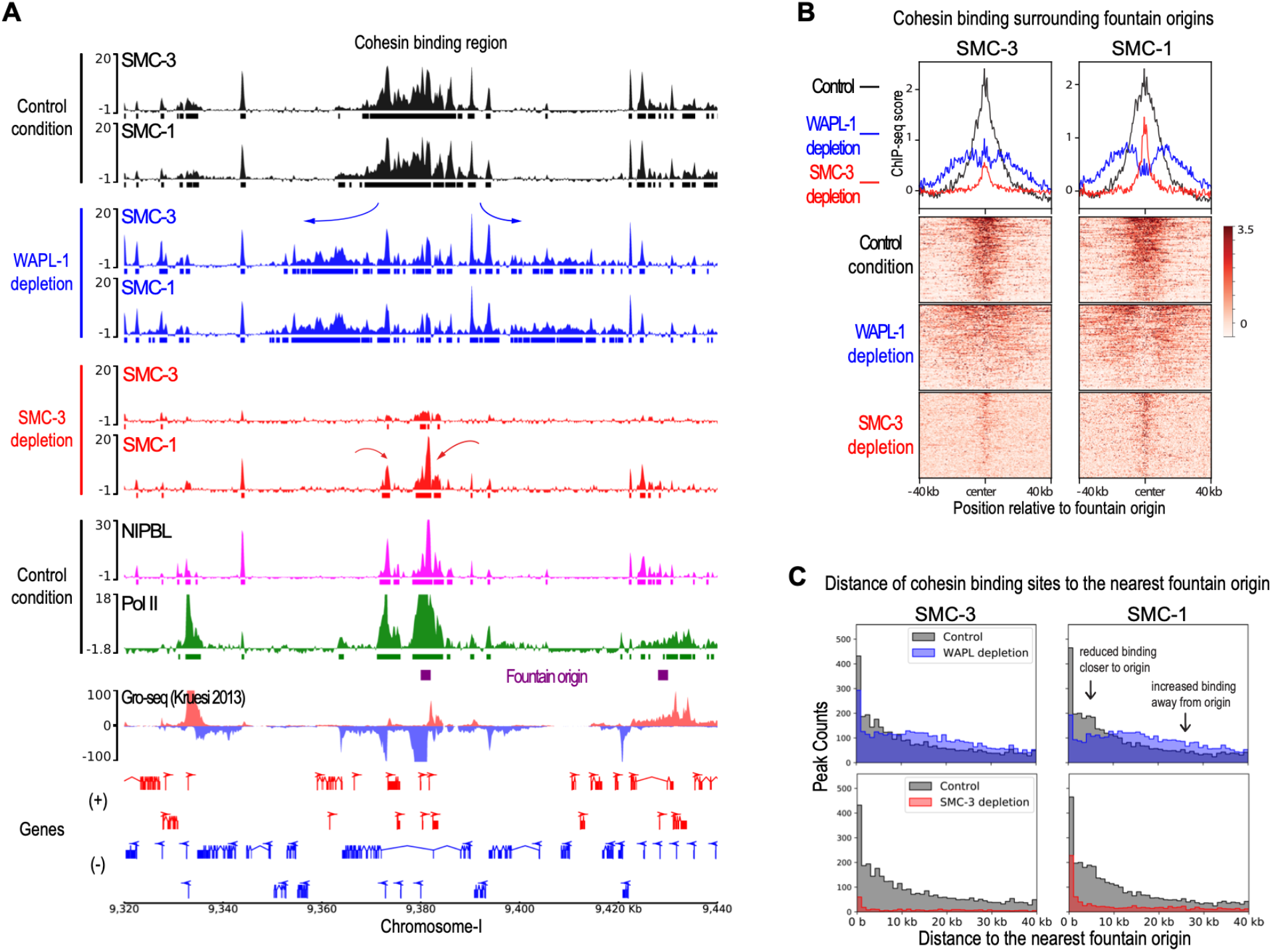
Cohesin spreads from fountain origins. A) Genome browser view around fountain origins. Plotted are input-subtracted ChIP-seq tracks for cohesin subunits, SMC-3 and SMC-1, in three conditions: control (black), WAPL-1 depletion (blue), and SMC-3 depletion (red). The ticks below signal tracks indicate MACS2 called peaks. Fountain origins are shown in purple. B) Average profile and heatmap of SMC-3 and SMC-1 with respect to fountain origins. Plotted are input-subtracted ChIP-seq tracks for cohesin subunits, SMC-3 and SMC-1, in three conditions: control (black), WAPL-1 depletion (blue), and SMC-3 depletion (red). C) Histogram of distance between cohesin subunit summit and the nearest fountain origin. The three conditions are control (gray), WAPL-1 depletion (blue), and SMC-3 depletion (red). Change in cohesin binding sites upon WAPL depletion is consistent between SMC-1 and SMC-3 and is highlighted in the panel corresponding to SMC-1.

Upon WAPL depletion, the cohesin binding region widens, reaching a span of nearly 40-kb (Figure 4A, Supplemental Fig S2, blue tracks). The increase in cohesin spreading upon WAPL depletion is reproducible (Supplemental Fig S3A) and generalizable across the fountain origins as measured by intensity (Figure 4B) or by binding site distance to the fountain origins (Figure 4C). This demonstrates that depletion of WAPL results in cohesin moving farther away from the fountain origins. If cohesins were moving towards the fountain origins, WAPL depletion would increase cohesin binding at the fountain origin. In contrast, WAPL depletion results in relatively lower cohesin at the fountain origins (Supplemental Fig S3C). Thus, we conclude that cohesins that are loaded at the fountain origins are moving away from the loading site in both directions, consistent with preferential loading followed by bi-directional loop extrusion implied by the Hi-C data (Figure 3C).

### Fountain origins harbor multiple NIPBL binding sites

NIPBL protein is important for cohesin recruitment to DNA (Ciosk et al., 2000). Thus, we next analyzed the distribution and function of the *C. elegans* ortholog of NIPBL, encoded by *pqn-85/scc-2* (Lightfoot et al., 2011). The strength of fountains positively correlated with both average NIPBL ChIP-seq signal (Figure 5A, left panel) as well as the number of NIPBL peaks (Figure 5A, right panel). Recent studies suggested that NIPBL is a processivity factor for cohesin loop extrusion (Alonso-Gil et al., 2023; Banigan et al., 2023; Busslinger et al., 2017; Ciosk et al., 2000; Han et al., 2023; Kagey et al., 2010). These models suggested that NIPBL is carried by cohesin to non-loading sites through loop extrusion activity (Han et al., 2023; Petela et al., 2018). We tested this prediction for *C. elegans* by performing ChIP-seq for NIPBL upon SMC-3 and WAPL depletion.

**Figure 5.**
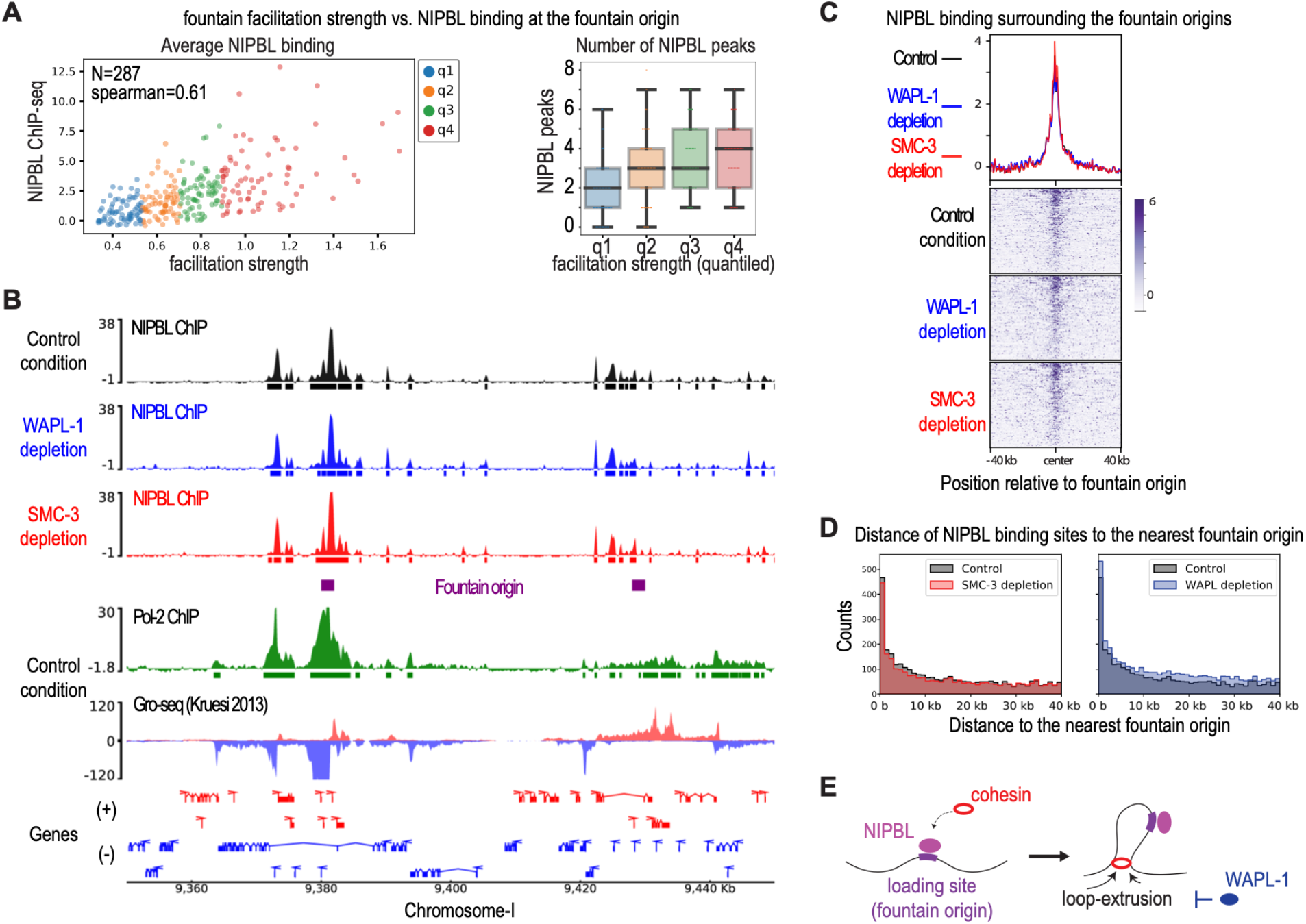
NIPBL correlates with fountain strength and its binding remains unchanged upon SMC-3 and WAPL-1 depletion. A) Left panel: Scatterplot of fountain facilitation strength vs. mean NIPBL ChIP-seq enrichment in control condition. Fountains divided into quartiles based on their strength are shown in different colors. Right panel: Boxplot of fountain facilitation strength (divided into quantiles) and the number of NIPBL summits within 6kb region. B) Genome browser view around fountain origins. Plotted are input-subtracted ChIP-seq track for NIPBL in three conditions: control (black), WAPL-1 depletion (blue), and SMC-3 depletion (red). The ticks below signal tracks indicate MACS2 called peaks. Fountain origins are shown in purple. C) Average profile and heatmap of NIPBL with respect to fountain origins. Plotted are input-subtracted ChIP-seq track for cohesin subunits, SMC-3 and SMC-1, in three conditions: control (black), WAPL-1 depletion (blue), and SMC-3 depletion (red). D) Histogram of distance between NIPBL summit and the nearest fountain origin. The three conditions are control (gray), WAPL-1 depletion (blue), and SMC-3 depletion (red). E) Cartoon model of cohesin moving away from fountain origins without NIPBL as it performs two-sided loop extrusion.

### Cohesin and WAPL depletion does not affect NIPBL binding pattern measured by ChIP-seq

If cohesin mediates NIPBL binding, a portion of NIPBL ChIP-seq signal must be SMC-3 dependent. This prediction was not met. Unlike SMC-1, NIBPL binding surrounding the fountains did not reduce upon SMC-3 depletion (Figure 5BC, red tracks), and the position of NIPBL peaks relative to the fountain origins did not change (Figure 5D, left panel). NIPBL antibody was previously validated by western blot analysis upon immunoprecipitation and RNAi depletion (Kranz et al., 2013). To confirm that ChIP-seq signal is dependent on NIPBL protein, we degron tagged *pqn-85* endogenously, and validated knockdown by western blot analysis (Supplemental Fig S4AB). Binding of NIPBL and SMC-3 surrounding the fountain origins were consistently lower upon NIPBL depletion in the presence of auxin, while H4K20me1 did not change (Supplemental Fig S4C-E). Therefore, NIPBL ChIP-seq signal is dependent on NIPBL protein, validating our conclusion that SMC-3 depletion did not significantly reduce NIPBL binding at fountains.

A second prediction of NIPBL translocating with cohesin is that upon WAPL depletion, NIPBL binding should also increase outwards from the fountain origins. This prediction was also not met. Unlike SMC-3, the NIPBL binding region did not become wider when WAPL was depleted (Figure 5BC, blue tracks). We noticed variability in the level of WAPL depletion between biological replicates and accounted for it by comparing NIPBL and SMC-3 binding in the same ChIP extracts (Supplemental Fig S5A). While slightly more NIPBL peaks were detected upon WAPL depletion (Supplemental Fig S5B), the effect was global and did not result in relative increase in NIPBL away from the fountain origins (Figure 5D, Supplemental Fig S5C). Further categorizing cohesin binding sites into two groups, ones uniquely present in the control condition and in WAPL depletion condition, also shows that despite clear redistribution of cohesin, no such phenomenon was observed for NIPBL (Supplemental Fig S6A-D). Consistently, while cohesin binding decreased at the fountain origin upon WAPL depletion (Supplemental Fig S3C), NIPBL binding did not (Supplemental Fig S5D). Together, these results suggest that NIPBL does not translocate significantly with cohesin in *C. elegans* (Figure 5E)

### Fountain patterns are influenced by the distribution of cohesin loading sites

If focal loading of cohesin is the basis of fountains, distribution of NIPBL binding sites should correlate with the diversity of fountain patterns. Indeed, fountains with a broad base of interactions have a wide distribution of shorter NIPBL peaks near the fountain origin. For instance, a fountain with a broader NIPBL binding region contains a small TAD-like feature that is converted to a fountain at a farther distance away from the main diagonal (Figure 6A). Broad distribution of NIPBL sites close to the origin of this particular fountain is consistent with simulation of “broad loading” producing TAD-like structures (Guo et al., 2022). Thus, we infer that the small TAD-like feature is due to multiple cohesins colliding with each other, forming a randomly distributed cohesin stack. The extension into a fountain is likely due to cohesins at the two ends of the stack that are not constrained (Figure 6A, cartoon).

**Figure 6.**
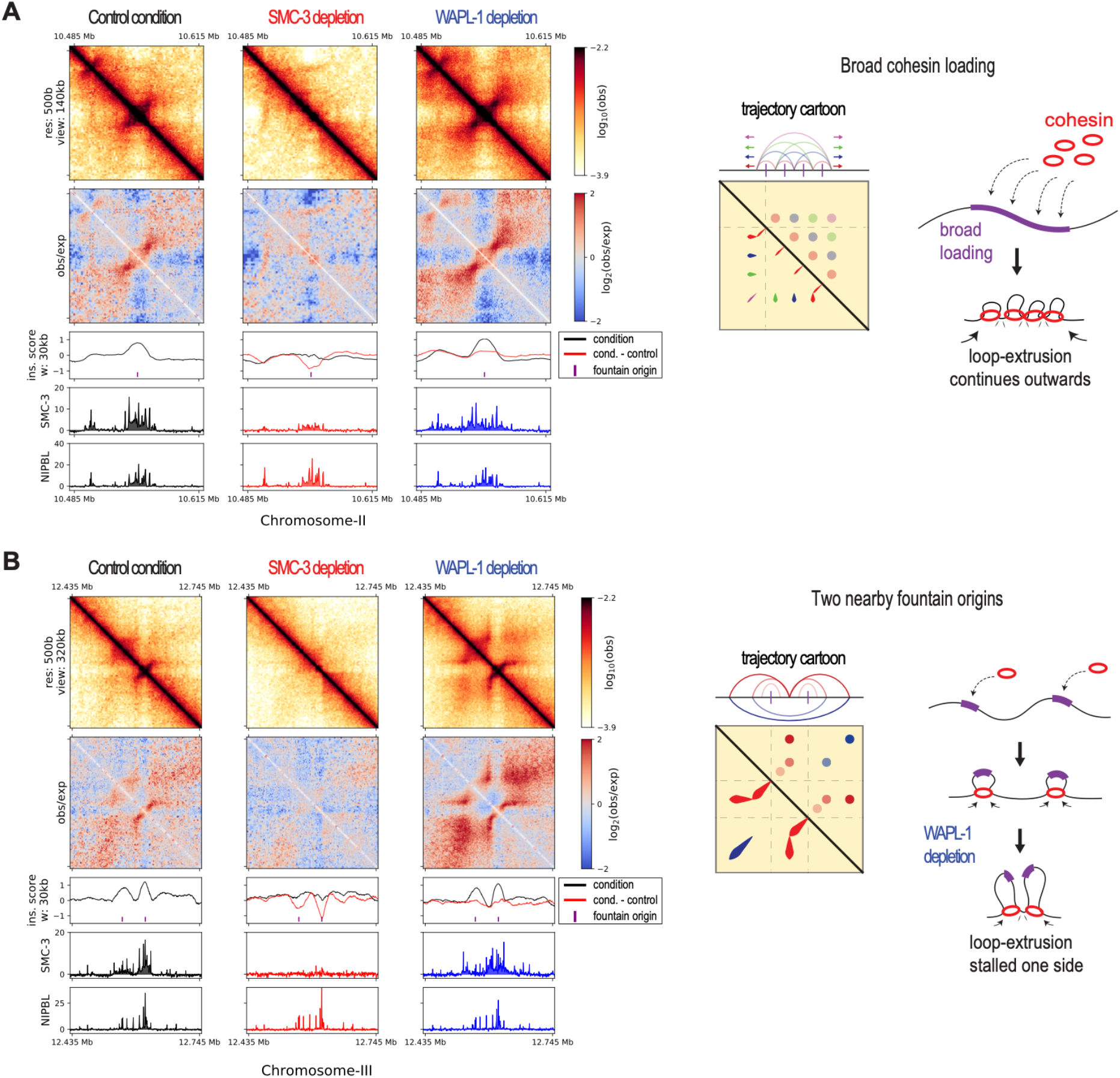
Fountains are heterogeneous. A) Hi-C snapshot of a locus containing broadly distributed NIPBL around fountain origins forming a TAD-like pattern. The trajectory cartoon shows one possible interpretation in which broadly loaded cohesins form multi-layered nested loops (blue, green, purple) giving rise to a broad region of enriched contact. B) Hi-C snapshot of a locus containing two nearby fountains forming ‘stripes.’ The trajectory cartoon shows one possible interpretation, where collision of two-sided loop extruding cohesins from two fountain origins stalling each other, causing them to behave as one-sided loop extruding factors (red). The collision results in nested loops, where the outer loop (blue) forms enriched contact away from the main diagonal.

### Diversity of fountain patterns support collision and stalling of cohesin molecules in vivo

The increased cohesin processivity upon WAPL depletion revealed possible collision between cohesin molecules. For example, two distinct fountains in the control condition extend to generate broad stripes upon WAPL depletion (Figure 6B). The stripes can emerge from cohesin coming from the left fountain origin stalled by the cohesin coming from the right fountain origin, and vice versa. Alternatively, a bidirectional barrier between the two fountain origins stalls cohesins loaded at either fountain. For this particular region, we favor cohesin collision, because the pronounced contact probability that appears off-diagonal suggests presence of cohesin-cohesin nested structure (Figure 6B, cartoon). This interpretation is supported by similar Hi-C patterns produced by simulation of two-sided loop extruding factors from two loading sites with low bypass rate (Brandao et al., 2021).

In simulation, preferentially loaded cohesins at fountain origins bumping into other or randomly distributed ‘background’ factors produce the fanning/dispersion property of the fountains (A. Galitsyna et al., 2023). Similarly, the loop-stacking model derived from *in vivo* imaging was proposed to emerge from collision among cohesins (Hafner et al., 2023). Our example fountain trajectories also suggest cohesin collides with itself or other chromatin associated factors.

Additional examples illustrate the heterogeneity of fountain patterns. For instance, one fountain shows a relatively narrow NIPBL binding and simple fountain trajectory (Supplemental Figure S7A), while another contains multiple NIPBL binding sites at the origin and along with more complex Hi-C interactions and fountain trajectory (Supplemental Fig S7B). In one example, we observe a mid-shift in the fountain’s trajectory, suggesting that there is an uneven distribution of factors that regulate cohesin movement (Supplemental Fig S7C). Lastly, we observe a ‘stripe-like’ feature embedded within a fountain (Supplemental Fig S7D), which could imply presence of a ‘stalling element’ or a special case of loading event (Arnould et al., 2021; Han et al., 2023). We surmise that the diverse fountain shapes emerge from heterogenous NIPBL/loading site distribution and other *in vivo* barriers/facilitators of loop extrusion.

### Fountain origins are close to active enhancers

To identify genomic elements that underlie the preferential loading of cohesins, we utilized genome annotations based on chromHMM from two previous studies (Daugherty et al., 2017; Evans et al., 2016). Fountain origins are closest to genomic regions designated as active enhancers compared to other annotations (Figure 7A, Supplemental Fig S8A). Because the nearest enhancer-promoter pairs have a median distance of less than 2 kb in *C. elegans* (Supplemental Fig S8B), fountain origins are often found in regions of high enhancer and promoter density (Supplemental Fig S9). At a region where they are sufficiently distant, fountain origin was closer to the strong ATAC-seq peak associated with the enhancer (Figure 7B, Supplemental Fig S8C). Thus, preferential loading of cohesin forming the fountains occurs at sites close to active enhancers.

**Figure 7.**
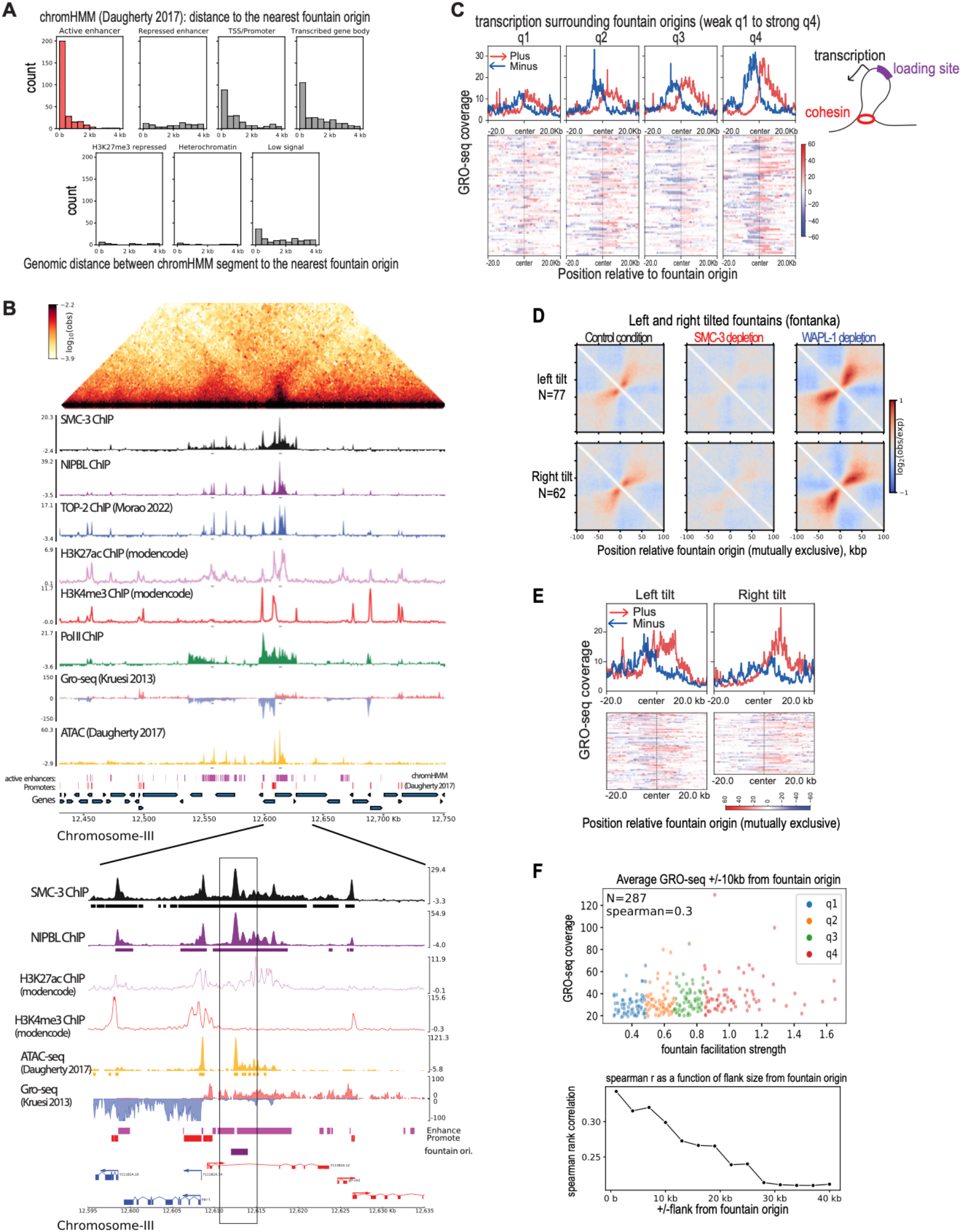
Fountain origins are close to active enhancers. A) Histogram of genomic distance between identified fountain origins in this paper and the annotated regions of chromHMM in (Daugherty et al., 2017). Active enhancers are highlighted in red. B) Genome browser snapshot of Hi-C and other transcription related features such as topoisomerase-II, H3K27ac, H3K4me3, Pol II, Gro-seq, and ATAC-seq (top panel). Zoom-in near a fountain origin is shown in the bottom panel. C) Average profile of Gro-seq data from (Kruesi et al., 2013) split into plus (red) and minus (blue) strands are plotted with respect to the fountain origins. The fountains are grouped into four quantiles based on their strength. On the right, the cartoon highlights that origins of strong fountains are often close to upstream of transcription. D) Left and right-directed fountains were identified using the fontanka tool (Galitsyna et al., 2023). The fountain origins of the two groups are mutually exclusive. E) GRO-seq plot centered around the left and right-directed fontanka fountain origins. F) Scatterplot showing relationship between facilitation strength of the fountain origin and the signal mean of GRO-seq (Kruesi et al., 2013) in 20 kb region centered at the fountain origin (top panel). Sweep of spearman correlation as a function of flanking region used to average GRO-seq signal (bottom panel).

Plotting published GRO-seq data (Kruesi et al., 2013) suggests that fountain origins are located upstream of transcribed regions (Figure 7C). To test if the direction of transcription correlates with trajectory of the fountains, we categorized them as left and right-directed using the fontanka tool (Figure 7D, Supplemental Figure 10A-C) (A. Galitsyna et al., 2023). While there was some bias in the distribution of Pol II binding sites with respect to left or right tilted fountains (Supplemental Fig S10D), correlation between the tilt and the direction of transcription was not strong (Figure 7E). Nevertheless, three lines of analyses, including example fountains coinciding with histone modifications and proteins associated with active transcription (Figure 7B, Supplemental Fig S8C), GRO-seq enrichment being lower around weak fountains and higher over strong ones (Figure 7C), and a positive correlation between transcription surrounding a fountain origin and fountain facilitation score (Supplemental Fig S8D) together demonstrate that fountains are associated with regions of active transcription.

### One hour depletion of SMC-3 or WAPL does not lead to changes in RNA Pol II binding

We used ChIP-seq analyses of RNA Pol II to analyze the effect of SMC-3 and WAPL knockdown. Unlike the striking changes observed in Hi-C, cohesin and WAPL depletion did not result in significant change in RNA Pol II distribution (Figure 8A). The lack of change in RNA Pol II ChIP-seq was reproducible (Supplemental Fig S11A) and generalizable across fountain origins (Figure 8B). Next, we categorized genes into two groups: one with higher cohesin binding in control and other upon WAPL depletion (Figure 8C). Despite the noticeable shift in cohesin binding between the two groups of genes upon WAPL depletion, no such change occurred for RNA Pol II (Figure 8D, Supplemental Fig S6E). Orthogonally, we observe that mRNA-seq showed minimal change upon 1 hour depletion of SMC-3 and WAPL (Figure 8E). We considered the possibility that chromatin may require more time to reach a new-found equilibrium upon loop extrusion perturbation (Nuebler et al., 2018). However, increasing the duration of auxin treatment to 4-hours also showed minimal change in mRNA-seq (Supplemental Fig S11B). Therefore, we conclude that acute depletion of cohesin and WAPL immediately and dramatically change cohesin binding and the 3D DNA contacts measured by Hi-C, but do not result in equally strong changes in gene expression.

**Figure 8.**
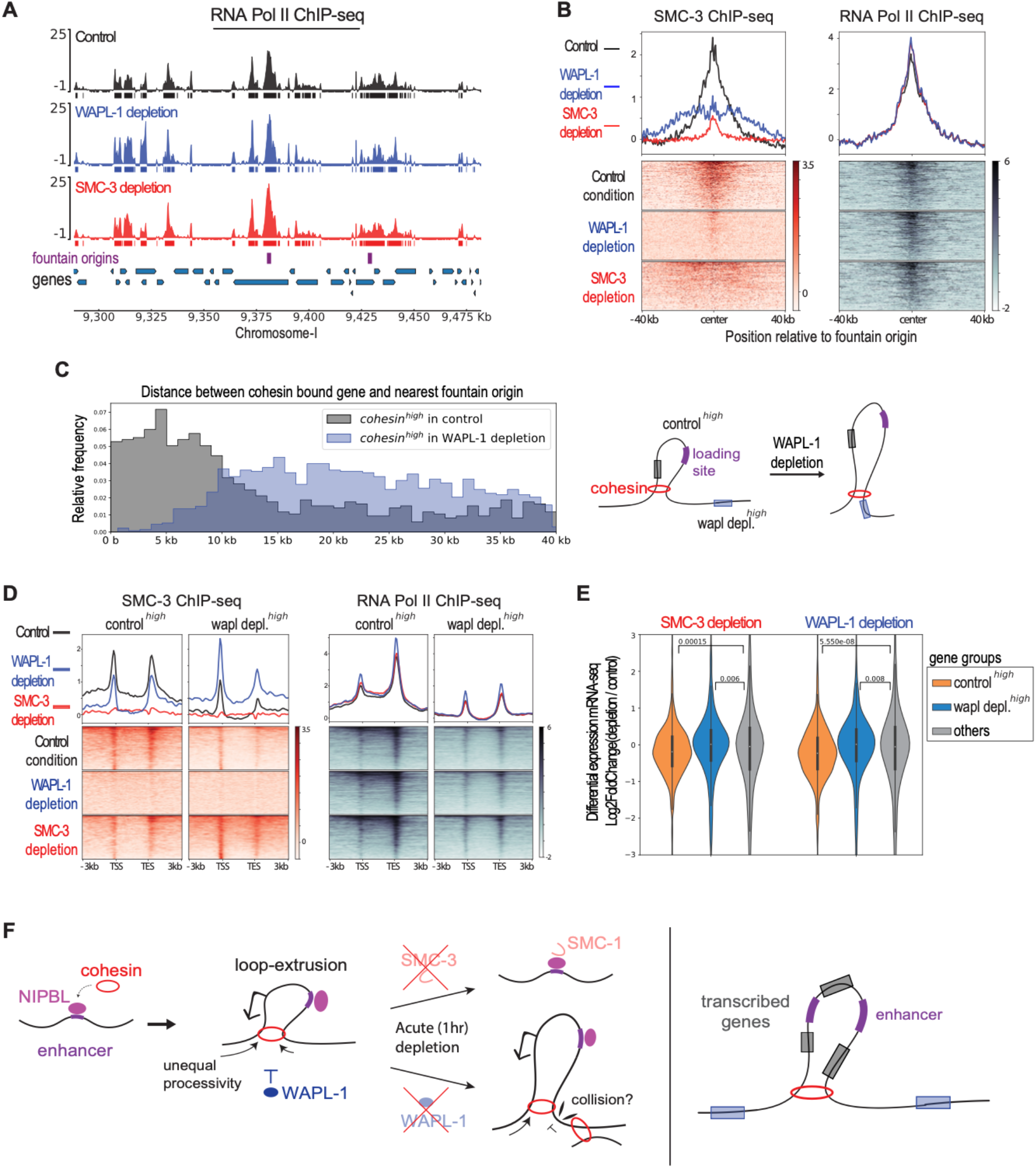
One hour depletion of SMC-3 or WAPL does not lead to drastic changes in RNA Pol II binding. A) Genome browser view around fountain origins. Plotted are input-subtracted ChIP-seq track for Pol II in three conditions: control (black), WAPL-1 depletion (blue), and SMC-3 depletion (red). The ticks below signal tracks indicate MACS2 called peaks. Fountain origins are shown in purple. B) Average profile and heatmap of SMC-3 and Pol II binding with respect to fountain origins. Plotted are input-subtracted ChIP-seq track SMC-3 and Pol II in three conditions: control (black), WAPL-1 depletion (blue), and SMC-3 depletion (red). C) Genes are categorized into cohesin high in control condition (gray) or in WAPL depletion condition (blue), and their relative positions are plotted as histogram from the nearest fountain origin. The two groups are mutually exclusive. The cartoon highlights that genes with higher cohesin binding in control are closer to fountain origins than genes with higher cohesin binding in WAPL depletion condition. D) Average profile and heatmap of SMC-3 and Pol II binding with over two groups of genes shown in C. Plotted are input-subtracted ChIP-seq track SMC-3 and Pol II in three conditions: control (black), WAPL-1 depletion (blue), and SMC-3 depletion (red). E) Differential mRNA-seq expression values (log2fc from DESeq2) is plotted for control high (n=817), wapl high (n=1268) genes that were further subsetted to be within 40 kb from the fountain origin, where the differential cohesin binding is the most pronounced. Other genes (n=15800) were used as control. Mann-Whitney test was used to generate p values. F) The proposed model of cohesin loading and loop extrusion events inferred from the perturbation experiments and literature. Activated enhancers are bound by NIPBL which increase the frequency of cohesin loading at or close to these sites. After loading, cohesin extrudes DNA in an effectively two-sided manner. Note that despite being depicted as a singular ring, the data cannot distinguish how many cohesin complexes are at the base of the loop. Upon acute depletion of SMC-3, the binding of NIPBL is unaffected and residual amount of SMC-1 is found at loading site. The extruded loop is abolished. Upon acute depletion of the negative regulator WAPL-1, cohesin further extrudes DNA in two-sided manner away from the jet origin. The extended extrusion revealed that the clash between incoming and outgoing cohesin molecules can stall extrusion. Together, cohesin processivity at each side can be unequal and may be facilitated by transcription moving in the same direction. The extruded loops can encompass multiple cis-regulatory elements and genes.

## Discussion

Cohesin and CTCF mediated TAD formation contribute to proper development and normal physiology in mammals, but many organisms, including yeast, worms and plants, contain cohesin but lack CTCF (Dong et al., 2017; Heger et al., 2012). Here, we used *C. elegans* as a model to address the question: what does interphase cohesin do in organisms without CTCF? We reasoned that the answer would point to the ancestral function of cohesin in gene regulation. Our approach was to apply the auxin-inducible degradation system to acutely deplete the cohesin ATPase subunit SMC-3 or its negative regulator WAPL-1 in somatic cells in *C. elegans* larvae. Our results are consistent with a model that preferential loading of cohesin followed by bi-directional loop extrusion organizes the 3D DNA contacts surrounding active enhancers (Figure 8F).

### Fountains are a conserved feature of 3D genome organization mediated by cohesin

Recent studies including this one demonstrate that cohesin mediated loops originating from distinct sites are the basis of Hi-C defined features termed jets, plumes, or fountains in model species of fungi, worms, zebrafish and mammals (A. Galitsyna et al., 2023; Guo et al., 2022; Isiaka et al., 2023; Ning Qing Liu et al., 2021; Shao et al., 2024; Wike et al., 2021). In zebrafish, fountains coincide with enhancer activity across development, and knockdown of pioneer transcription factors eliminate fountains originating at these sites (A. Galitsyna et al., 2023). In mammalian cells, analysis of cohesin binding upon CTCF site deletion, depletion of pioneer transcription factors, and ectopic recruitment experiments suggest that cohesin is preferentially loaded at enhancers (Dowen et al., 2013; Han et al., 2023; N. Q. Liu et al., 2021; Vos et al., 2021). Our work also showed an overlap of fountain origins with active enhancers in *C. elegans*, thus preferential loading of cohesin may be an evolutionarily conserved mechanism of regulating 3D DNA contacts surrounding enhancers.

### NIPBL recruitment to active enhancers may facilitate cohesin loading

Based on analysis of individual fountains and their genomic context, we observed that the strength and the pattern of 3D DNA contacts forming the fountains correlates with NIPBL binding. Immunofluorescence analyses of cohesin in the germline showed that *C. elegans* NIPBL (PQN-85/SCC-2) is required for cohesin binding to chromosomes (Ciosk et al., 2000; Lightfoot et al., 2011; Seitan et al., 2006). We also observed lower cohesin binding upon NIPBL depletion (Supplemental Figure S4). In many organisms, including *C. elegans*, NIPBL binds to active enhancers (Busslinger et al., 2017; Kranz et al., 2013; Zhu et al., 2021). In yeast, NIPBL interacts with and is recruited by the mediator complex (Mattingly et al., 2022). Therefore, it is possible that transcription factors interaction with the mediator recruits NIPBL to load cohesin at active enhancers.

In mammals, NIPBL is proposed to be a processivity factor that regulates the engagement of cohesin with DNA during loop extrusion (Alonso-Gil et al., 2023). ChIP-seq analysis of NIPBL upon RNAi depletion of a cohesin subunit in HeLa cells suggested that NIPBL binding at promoters is cohesin dependent (Banigan et al., 2023). Ectopic loading of cohesin by TetO-tetR system showed that NIPBL binding spreads along with cohesin, suggesting that cohesin carries NIPBL as it moves (Han et al., 2023). In *C. elegans*, we did not observe a change in NIPBL binding upon cohesin or WAPL depletion (Figure 5). Therefore, it is unlikely that a significant amount of NIPBL protein translocates along with loop extruding cohesin in *C. elegans*. It is possible that the longer processivity of mammalian cohesin compared to that of *C. elegans* is related to NIPBL’s ability to translocate with cohesin in a larger genome. Differences between cohesin and its regulators in mammals versus *C. elegans* may provide insights into the conserved and distinct mechanisms during the evolution of animal genomes.

### Collision of loop extruding cohesins creating a barrier effect

Modeling approaches are used to connect the behavior of single cohesin molecules to the population average data produced in Hi-C and we have interpreted our data in light of these studies (Banigan et al., 2020). Our results suggest that frequent loading results in the collision of loop extruding cohesin molecules when these molecules are not removed from the DNA template by WAPL (Figure 6). The resulting structure suggests cohesin continues to loop extrude on one side, while being stalled on the other. Therefore, an asymmetry in the loop extrusion is introduced by a barrier, in this case an incoming cohesin. Supporting our interpretation, computational modeling used the interaction of outgoing and incoming cohesin molecules to explain the dispersion property of fountains in zebrafish (A. Galitsyna et al., 2023), which are similar to the fountains in *C. elegans* and unlike the hairpin structures formed by preferential loading and extrusion by bacterial SMCs (Brandao et al., 2021).

Collision between loop extruders may be a general feature of SMC complexes. In *C. elegans*, a specialized condensin I complex for dosage compensation (DC) is loaded at distinct *cis*-regulatory elements on the X chromosomes (Kim et al., 2022). The loading sites also form TAD boundaries by blocking loop extrusion by condensin DC (Anderson et al., 2019; Rowley et al., 2020). It is possible that the frequent loading of condensin DC at one recruitment site blocks the incoming molecules from another, because condensin DC loop extrudes in one direction (Kim et al., 2022). Intriguingly, we observed that cohesin-mediated loops are shorter on the X chromosome, raising the possibility that condensin DC may also stall cohesin, thus different SMC complexes may also collide.

### RNA Polymerase II: barrier or facilitator of cohesin mediated loop formation?

Our cohesin ChIP-seq data in *C. elegans* produced a pattern consistently observed in many organisms, where distinct peaks of binding are detected at promoters and enhancers (Peters et al., 2008; Rittenhouse & Dowen, 2024). We also observe that prominent fountains are formed at regions of high transcription. Earlier studies in yeast suggested that RNA Pol II may push cohesin (Glynn et al., 2004; Lengronne et al., 2004). More recent work used the framework of loop extrusion model and proposed that RNA Pol II and MCM complexes may be barriers to cohesin mediated loop extrusion (Banigan et al., 2023; Dequeker et al., 2022). It is possible that transcription facilitates the movement of cohesins at fountains. Alternatively, intermittent stalling by polymerase allows for better capturing of the loops by crosslinking-dependent methods like Hi-C and ChIP-seq. Facilitation or barrier activity of transcription may be based on the transcription machinery itself or the associated changes to DNA topology (Guerin et al., 2024; Morao et al., 2022). Future work is needed to determine if the association between transcription and fountains is causal or correlational due to sharing a common mechanistic origin.

### Fountains and gene regulation

In our study, one hour depletion of cohesin and WAPL resulted in striking changes in cohesin binding and the Hi-C contacts forming fountains, but not on RNA Pol II binding or mRNA levels measured by ChIP-seq and mRNA-seq, respectively (Figure 8). In a recent preprint, TEV-mediated cleavage of two cohesin kleisin subunits over a period of ∼24 hours from L1 larvae to L3 was used to demonstrate the role of COH-1 in fountain formation and address cohesin function in gene regulation in *C. elegans* (Isiaka et al., 2023). Authors reported a slight increase in the expression of genes closer to fountain origins upon cohesin cleavage. This contrasts with the slight decrease we observed (Figure 8E). In both studies, mRNA-seq was performed in whole animals and the effect was subtle. Thus, further research is required to address cohesin function in gene regulation.

In mammals, acute knockdown of cohesin also results in subtle changes in gene expression, yet mutations that reduce cohesin dosage or TAD boundaries produce significant changes (Cummings & Rowley, 2022; Hafner & Boettiger, 2023). It is possible that cohesin acts as a fine-tuner of transcription, or function earlier in development, and once an epigenetic state is established, short-term depletion of cohesin does not result in changes in transcription (Pelham-Webb et al., 2020).

### Evolution of cohesin activity

The co-evolution of underlying linear genome complexity and cohesin function may explain CTCF being lost in *C. elegans*. In mammals, the relatively large cohesin loops are constrained by positioning of CTCF to prevent ectopic activation of promoters (Hnisz et al., 2016; Levine et al., 2014). *C. elegans* cohesin makes ∼10 fold smaller loops compared to the mammalian cohesins, which seems to scale with gene density (∼one gene per 5 kb versus 100 kb). Our study offers two ideas for the shorter processivity of *C. elegans* cohesin. First in a compact genome, frequent encounters between incoming and outgoing cohesin molecules, as well as other SMC complexes and *in vivo* barriers may limit cohesin translocation. Second, intrinsic factors may also reduce processivity. For instance, NIPBL may have a more limited role in the processivity of cohesin in *C. elegans* compared to that in mammals.

Whether cohesin mediated DNA loops reduce *cis*-element search space (Hsieh et al., 2022), or functions to cyclically accumulate factors for promoter activation (Xiao et al., 2021), the relatively weak cohesin of *C. elegans* fits its underlying compact genome, where enhancers are intragenic or immediately upstream of promoter (Reinke et al., 2013). The presence of fountains in *C. elegans* argues that preferential loading of cohesin at active enhancers is an evolutionarily conserved mechanism of 3D genome regulation regardless of genome size and CTCF presence. Within the context of a compact animal genome, how cohesin-mediated organization of enhancer contacts contributes to cell-type specific gene expression is an intriguing question for future studies.

## Acknowledgements

JK and SE, and research in this manuscript were supported by NIGMS of the National Institutes of Health under award number R35 GM130311. Jun Kim was partially supported by NIGMS Predoctoral Fellowship T32HD007520. We thank Dominic Balcon for testing cohesin antibodies and Gencore at the NYU Center for Genomics and Systems Biology for sequencing and raw data processing.

## Author Contributions

JK and SE conceptualized the project and designed experiments. JK and HW performed experiments. JK performed data analysis. JK and SE prepared figures and wrote the manuscript.

## Supplemental

**Supplemental Figure S1.**
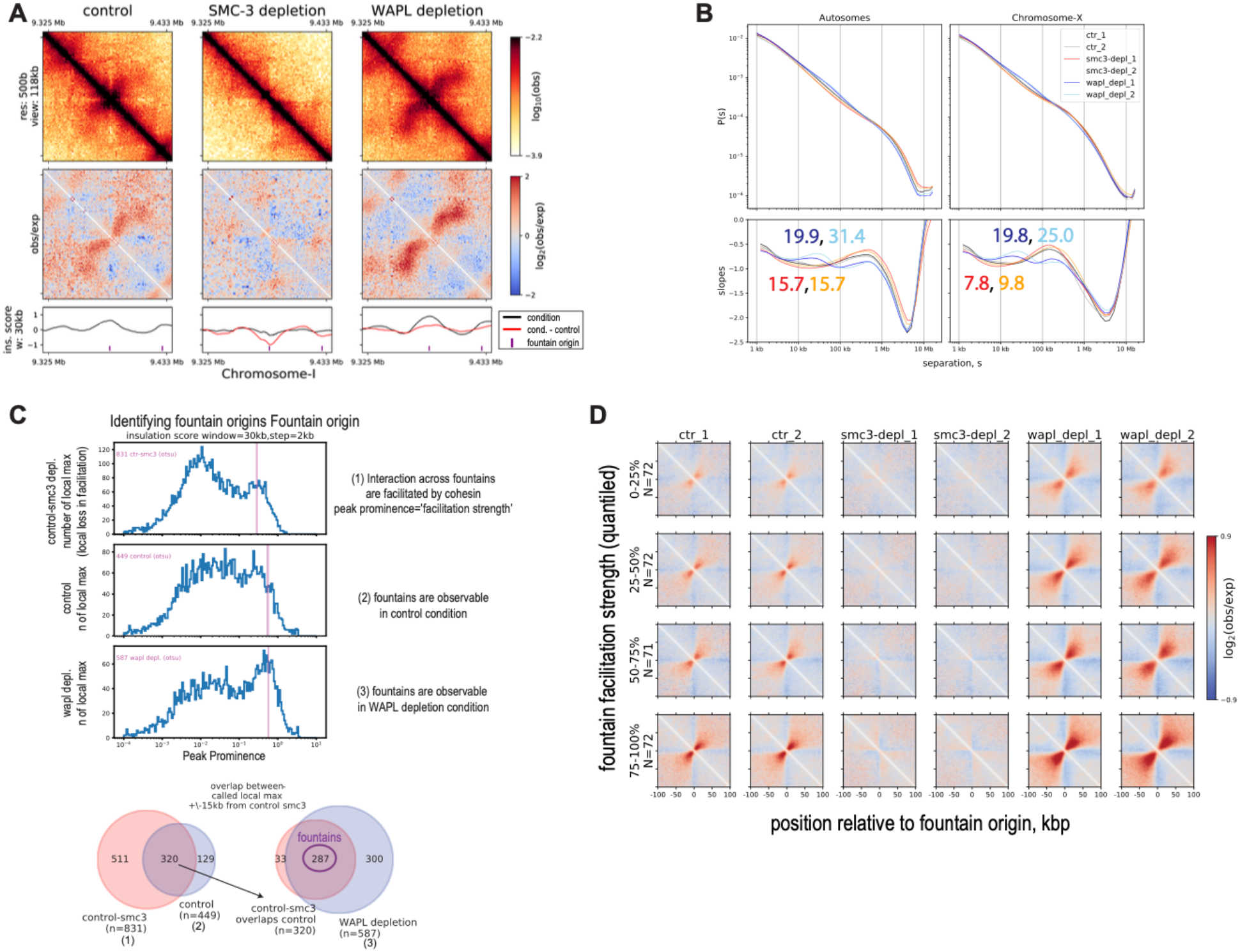
Fountain origin identification. A) Kinked fountain that begins as left-directed and then changes trajectory to the right. B) P(s) and log-derivative plots for three conditions separated by replicates. The numbers indicate inferred loop size of cohesin. C) Top panel: histogram of local maxima with varying degree of peak prominence. We define fountain to have the following properties: (1) interaction across the origin is facilitated by cohesin, (2) fountain is observable in control condition and (3) in WAPL depletion condition. These three conceptual criteria are applied computationally as identifying significant local maxima of (1) insulation scores delta (control condition minus SMC3 depletion condition), (2) insulation scores in control condition, and (3) insulation scores in WAPL depletion condition. The ‘facilitation strength’ corresponds to the peak prominence of insulation scores delta in (1), i.e. degree with which the interaction across the origin is facilitated by cohesin. The significant local maxima are called based on Otsu’s method of signal-noise separation indicated by vertical lines on the histogram. Bottom panel: pairwise overlap between significant local maxima called from criteria in A). Due to shift in the maxima occurring from tilt of the fountains, the ‘overlap’ allows for +/-15kb maximum distance (half of the insulation score window) from the local maxima called by insulation delta (1). We reason that insulation delta (1) provides the most accurate position and strength of the fountains since it corrects for underlying Hi-C structure that is not cohesin dependent. D) On-diagonal pileup of observed-over-expected matrix for each replicate for three conditions is centered at fountain origins divided into quantiles based on the ‘facilitation strength’.

**Supplemental Figure S2.**
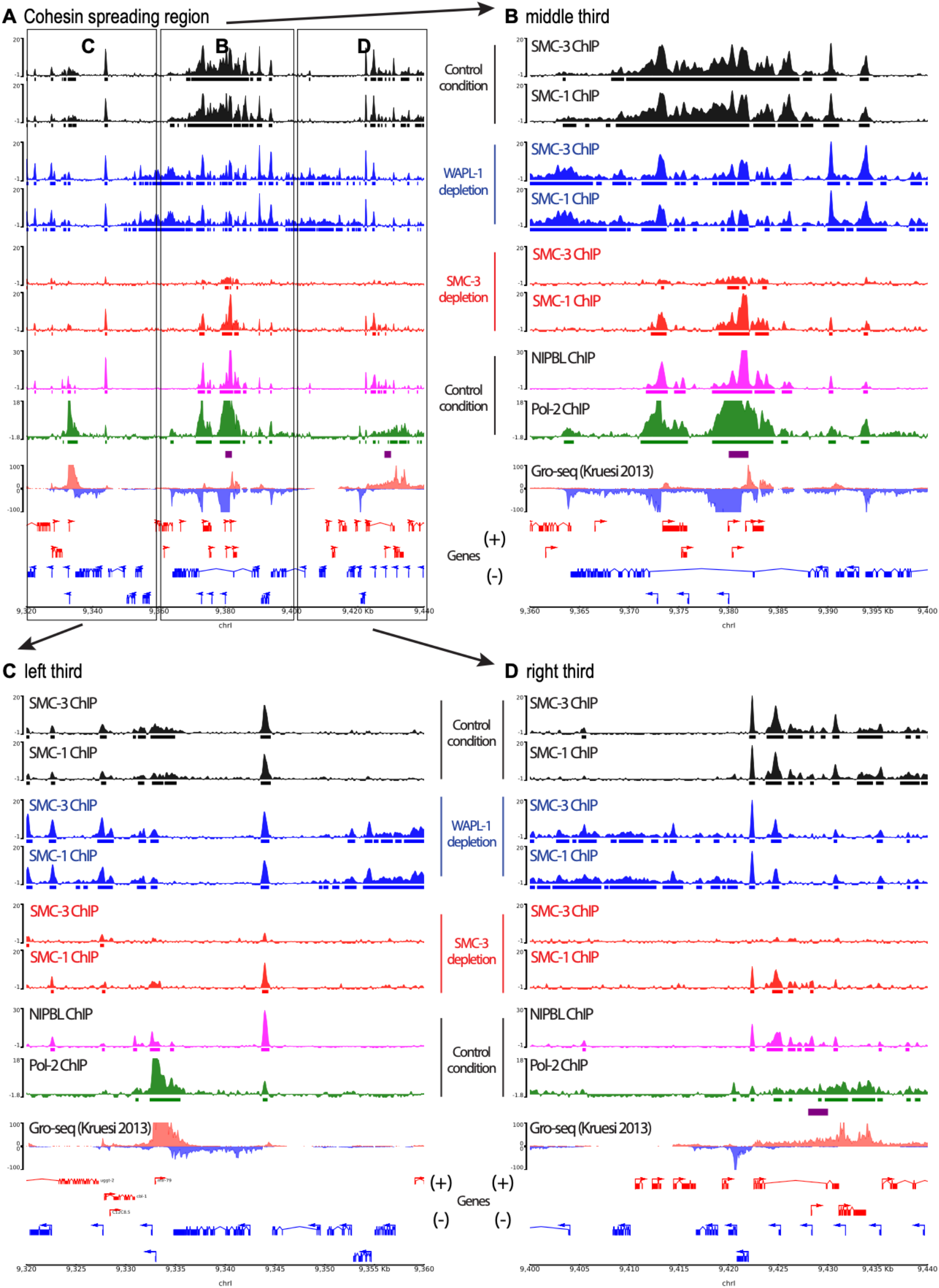
Zoom-in of spreading region of cohesin. A) A region of cohesin binding near fountain origins. The region is subdivided into three segments (B, C, D) to highlight changes in cohesin binding sites upon WAPL depletion. B) Zoom-in of the middle segment C) Zoom-in of the left segment D) Zoom-in of the right segment

**Supplemental Figure S3.**
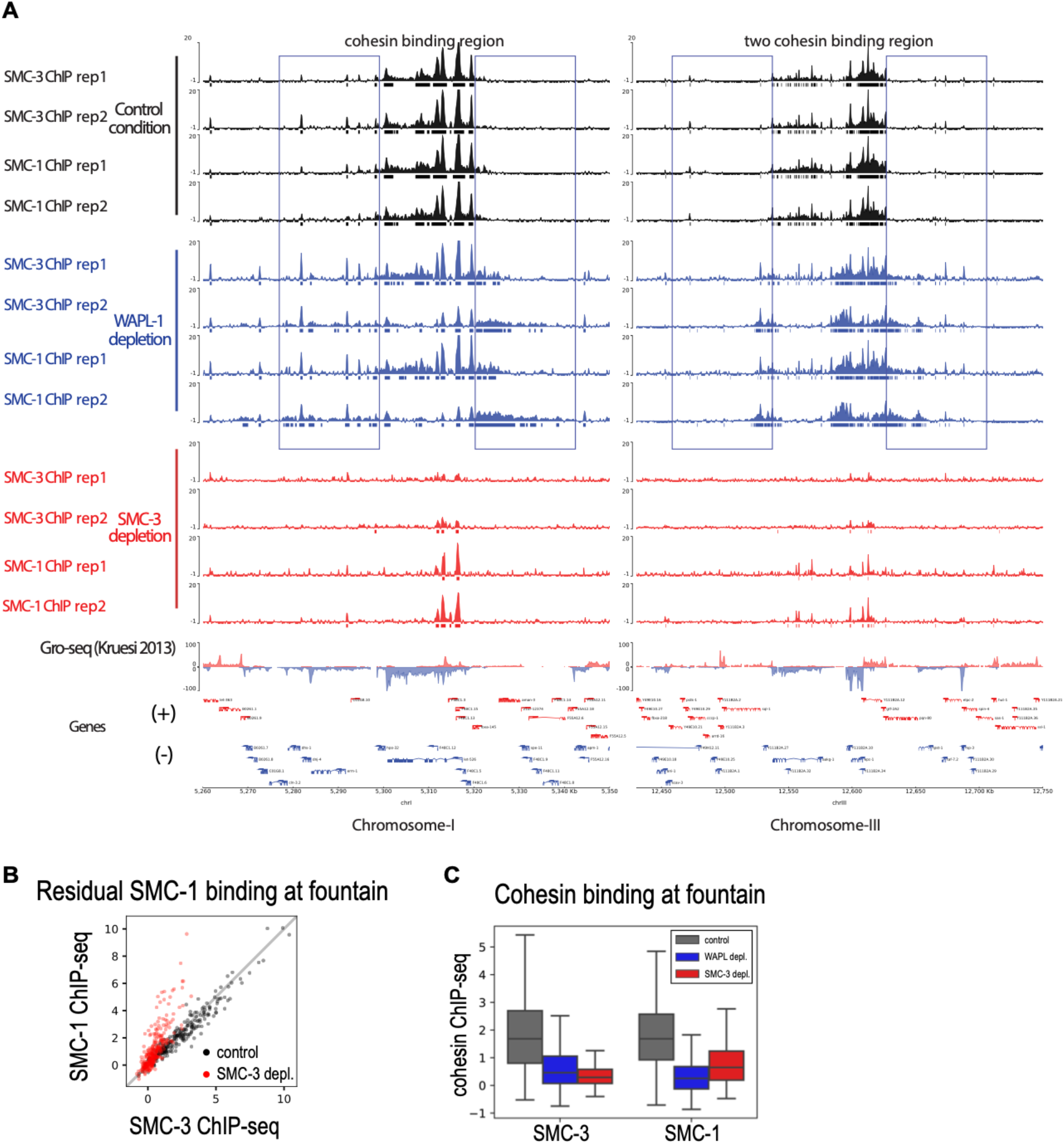
Replicates of SMC-1 and SMC-3 ChIP-seq data in control, SMC-3 and WAPL depletion conditions. A) Genome browser view of input subtracted ChIP replicates corresponding to representative regions. Rectangles are drawn to highlight the outward translocation of cohesin upon WAPL depletion. B) SMC-1 and SMC-3 signal averaged over 6kb region centered at fountain origins. SMC-1 and SMC-3 show similar binding across fountains origins in control condition, but this binding is SMC-1 skewed upon SMC-3 depletion. C) Boxplot of SMC-3 and SMC-1 binding at fountain origins. Input-subtracted ChIP-seq signal over 6kb region centered at fountain origin is averaged.

**Supplemental Figure S4.**
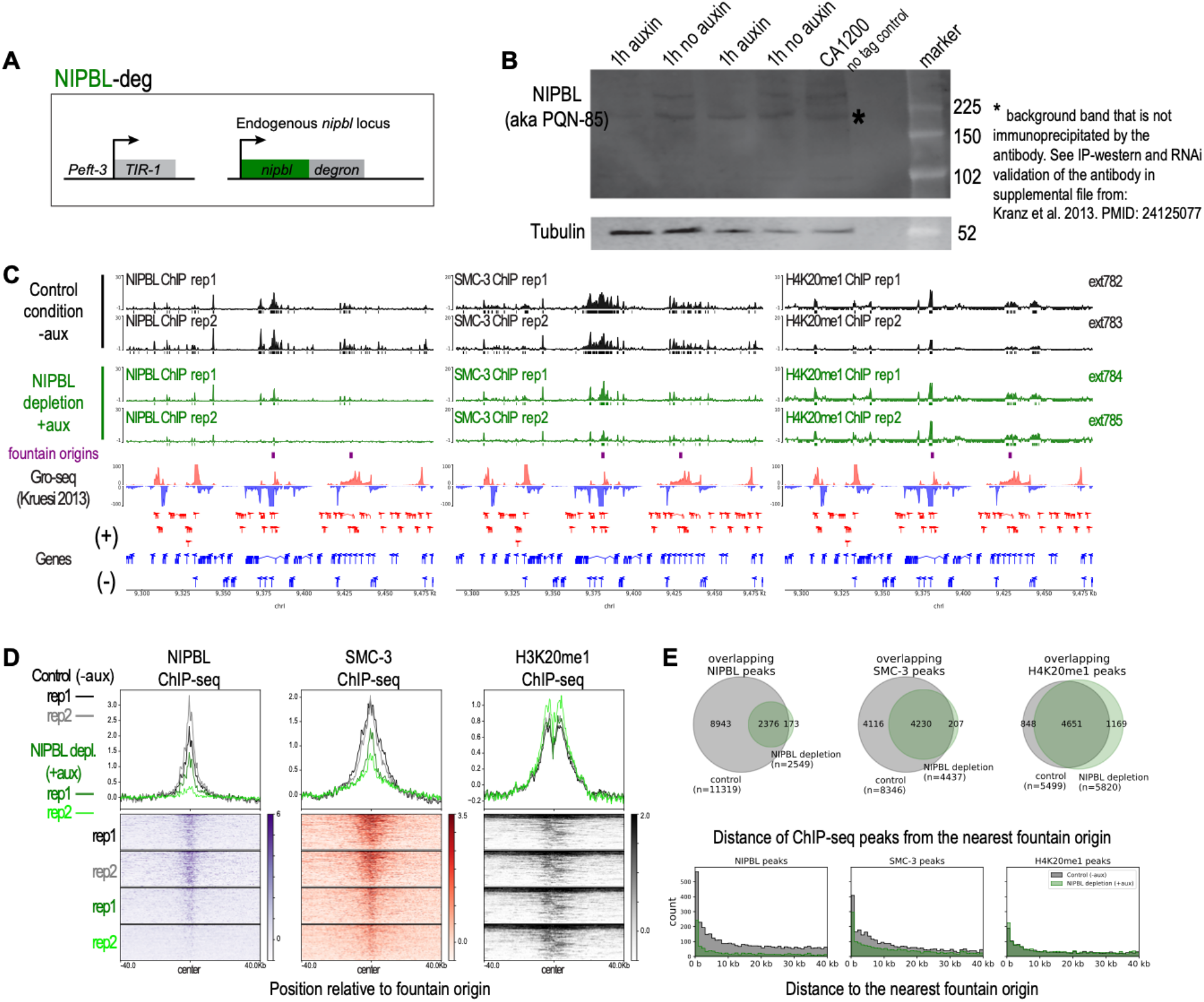
NIPBL depletion. A) Transgenic strain used in this figure. Exogenously inserted TIR1 is expressed in somatic cells using the eft-3 promoter. Endogenous nipbl (encoded by pqn-85/scc-2 gene) is C-terminally tagged with degron. B) Western blot images of L3 stage worms. CA1200 is TIR-1 control lacking the degron tag. Please note that the western blot replicates do not correspond to the ChIP extract replicates. This experiment was done to validate that NIPBL is knocked down in 1 hour with our transgenic strain. C) Genome browser view of ChIP replicates for the two conditions with (black) and without auxin (green). Two replicates of NIPBL (left panel), SMC-3 (center panel), and H4K20me1 (right panel) are shown. The same chromatin extract was used as indicated on the right label to reduce potential technical and biological variability. NIPBL and SMC-3 show depletion upon auxin treatment. H4K20me1 controls for the quality of the chromatin extract. D) Average profile and heatmap across fountain origins. Upon NIPBL depletion, both NIPBL and SMC-3 ChIP-seq signals reduce but not H4K20me1. Note that the saturated signal on top of the heat maps for H4K20me1 is due to X chromosome specific enrichment linked to dosage compensation (Albritton & Ercan, 2018). E) Venn-diagram of binding site changes between with and without auxin treatment (top panel). Histogram of binding site distribution relative to the fountain origins (bottom panel).

**Supplemental Figure S5.**
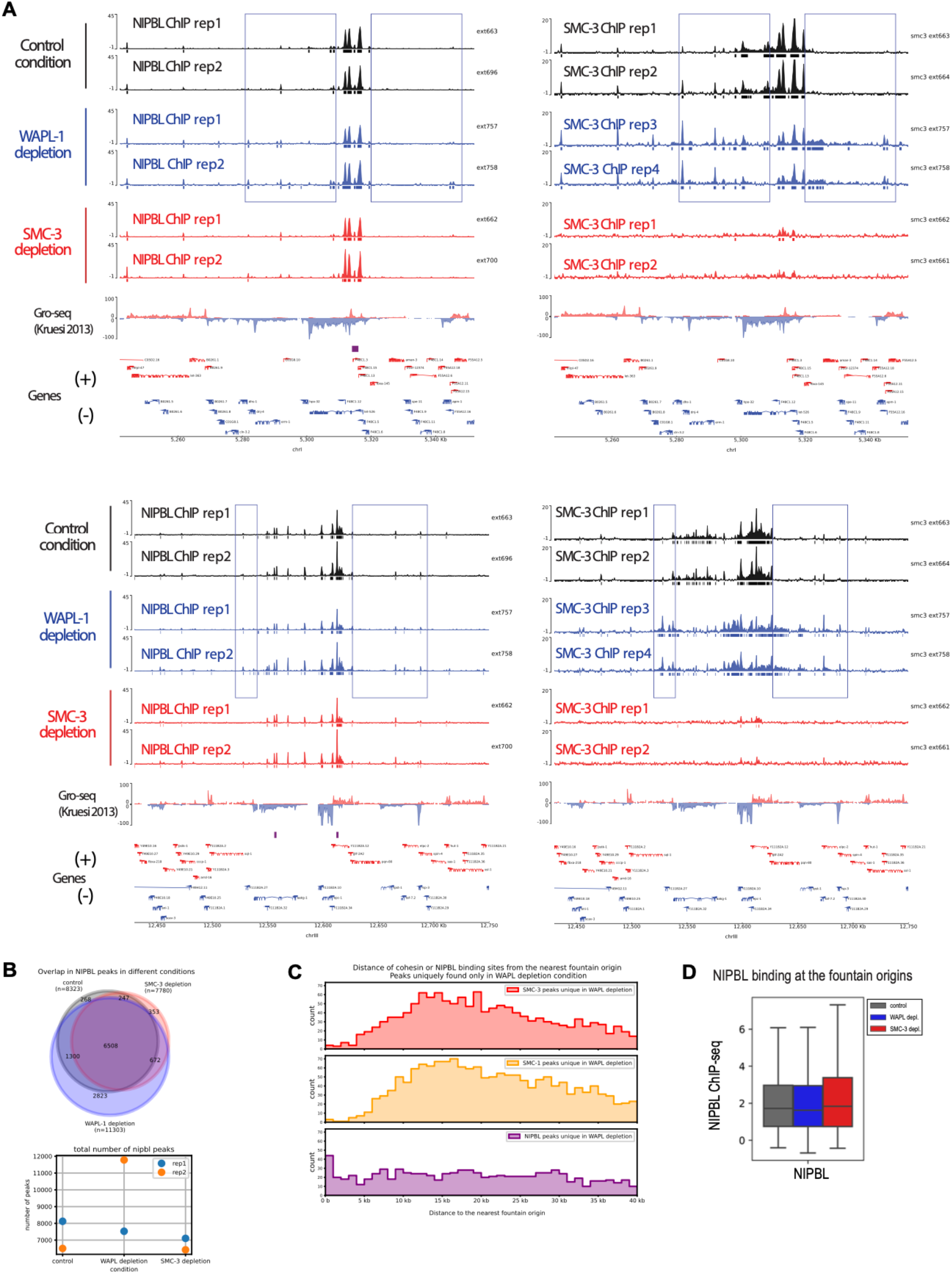
Replicates of NIPBL ChIP-seq data in control, SMC-3 and WAPL depletion conditions. A) Genome browser view of ChIP replicates. NIPBL ChIP-seq tracks for the control, WAPl-1, and SMC-3 depletion conditions (left) compared to the SMC-3 ChIP-seq tracks in the same conditions (Right). The ChIP extract identification numbers are provided to compare the effect of WAPL depletion on SMC-3 and NIPBL in the same ChIP extract and account for biological variability. The rectangles are used to highlight regions outside the cohesin binding regions in control where SMC-3 binding spreads out to in WAPL depletion. The same regions did not show increased NIPBL binding highlighting that unlike SMC-3, NIPBL localization is not dependent on WAPL. B) Venn-diagram of NIPBL peaks in three conditions (top). The total number of called peaks for each replicate (bottom). C) Histogram of SMC-3, SMC-1, and NIPBL ChIP-seq peak distribution with respect to the fountain origins. Only the peaks uniquely found in WAPL depletion are used. Unlike SMC-3 or SMC-1 peaks, the NIPBL peaks are not enriched away from the fountain origins, thus NIPBL did not translocate with cohesin upon WAPL depletion. D) Boxplot of NIPBL binding at fountain origins. Input-subtracted ChIP-seq signal over 6kb region centered at fountain origin is averaged. In contrast to SMC-3 or SMC-1 binding at fountain origins (Supplemental Fig S3C), NIPBL binding remains intact across perturbation.

**Supplemental Figure S6.**
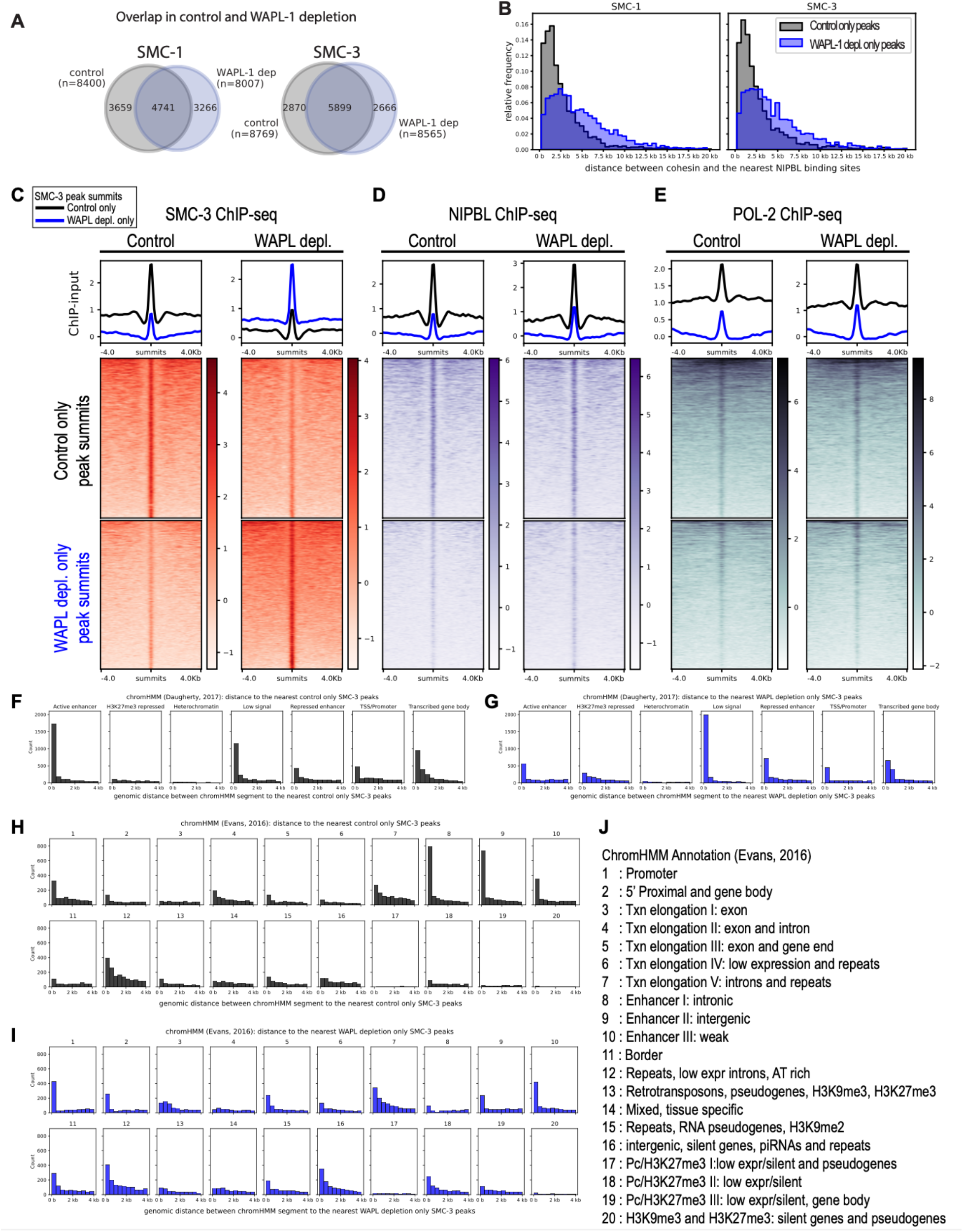
Investigation of differential cohesin binding sites. A) Venn-diagram showing overlap between cohesin binding sites between control and WAPL depletion conditions. B) non-overlapping peaks from A) are used to identify control specific peaks and WAPL depletion specific peaks. Two groups of peaks are used to plot the histogram of distance between cohesin binding site and NIPBL binding site in the corresponding condition. C) Average profile and heatmap of input subtracted SMC3 ChIP-seq track in control or WAPL-depletion conditions for two groups of SMC-3 summits from A). D) Average profile and heatmap of input subtracted NIPBL ChIP-seq track in control or WAPL-depletion conditions for two groups of SMC-3 summits from A). E) Average profile and heatmap of input subtracted Pol II ChIP-seq track in control or WAPL-depletion conditions for two groups of SMC-3 summits from A). F) Histogram of distance between control specific SMC-3 summits and annotated chromHMM in (Daugherty et al., 2017). G) Histogram of distance between WAPL depletion specific SMC-3 summits and annotated chromHMM in (Daugherty et al., 2017). H) Histogram of distance between control specific SMC-3 summits and annotated chromHMM in (Evans et al., 2016). I) Histogram of distance between WAPL depletion specific SMC-3 summits and annotated chromHMM in (Evans et al., 2016). J) Annotation for chromHMM in H,I. Taken from (Evans et al., 2016).

**Supplemental Figure S7.**
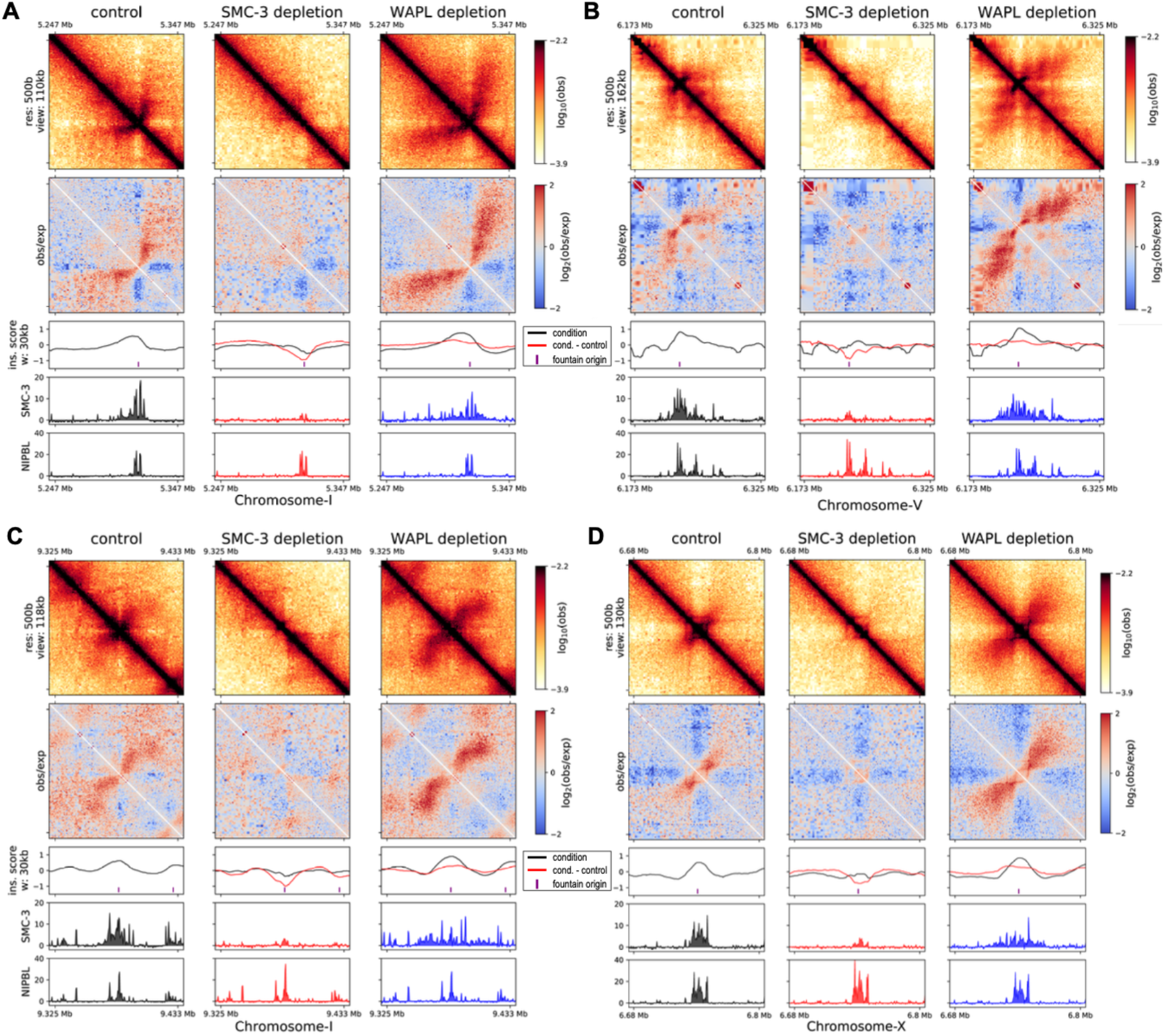
Heterogeneity of fountains. Hi-C plots of four regions with insulation scores, SMC-3, and NIPBL ChIP-seq data. A) Left-directed fountain. B) Right-directed fountain. C) Kinked fountain that begins as left-directed and then changes trajectory to the right. D) Fountain with embedded stripe pattern.

**Supplemental Figure S8.**
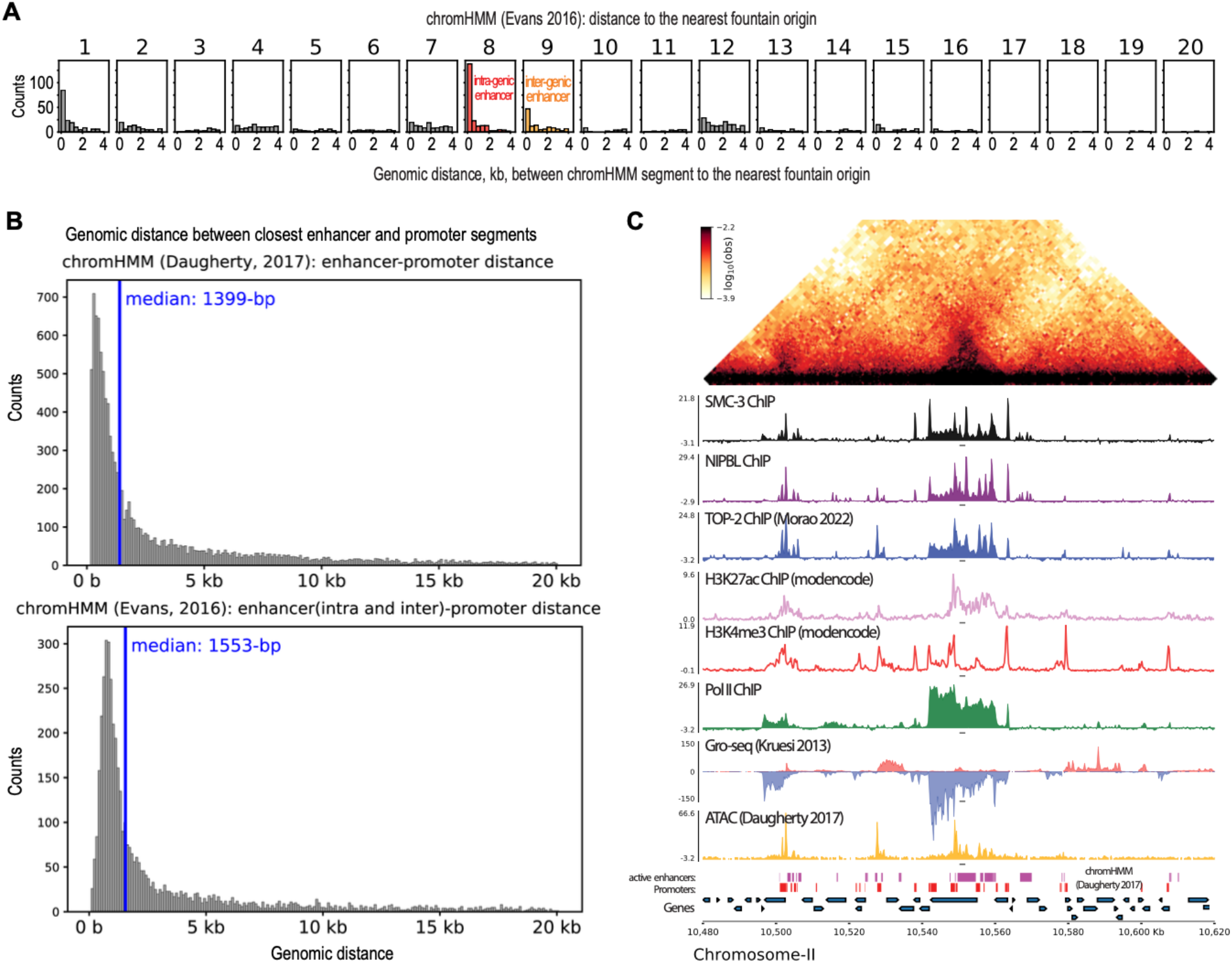
Fountains origins coincide with enhancers. A) Genome browser snapshot of Hi-C and other transcription related features such as topoisomerase-II, H3K27ac, H3K4me3, Pol II, Gro-seq, and ATAC-seq. B) Histogram of distance between enhancer and promoter based on chromHMM annotations. C) Histogram of 1kb binned genomic distance between identified jet origin in this paper and the annotated regions of chromHMM (Evans et al., 2016). Segments of chromHMM are subsampled to match the group containing the fewest number of segments; the shown histogram is an average of histograms generated from ten distinct random subsampling events. The two active enhancer groups are annotated. For detailed description of each group can be found in Supplemental Fig S6J. The annotations are directly taken from the original paper (Evans et al., 2016).

**Supplemental Figure S9.**
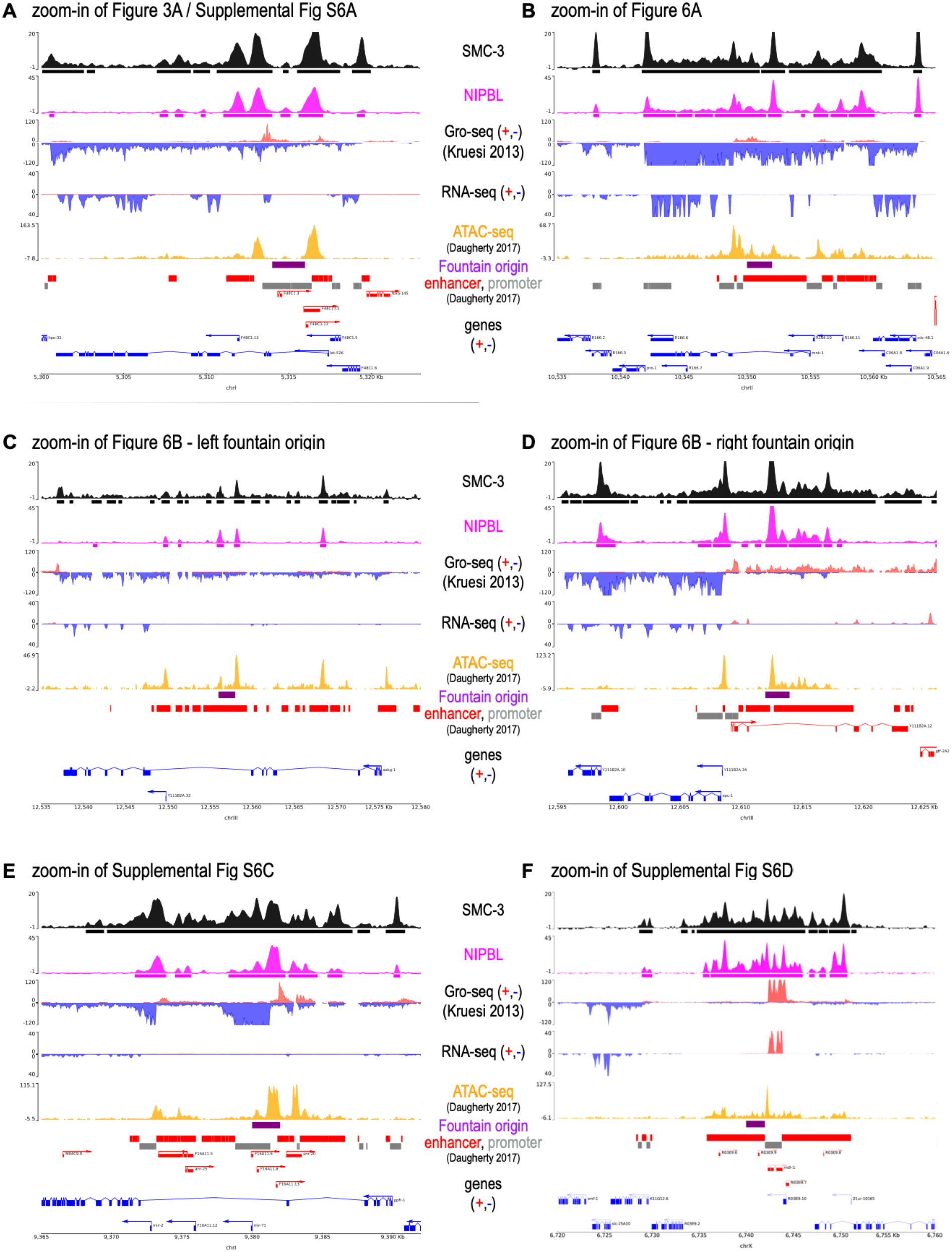
Zoom-in of various regions around fountain origins. The ChIP-seq tracks (SMC-3 and NIPBL) and RNA-seq track are from this study. Other key features such as Gro-seq (Kruesi et al., 2013) and ATAC-seq (Daugherty et al., 2017) are also plotted as well as the chromHMM from the same study.

**Supplemental Figure S10.**
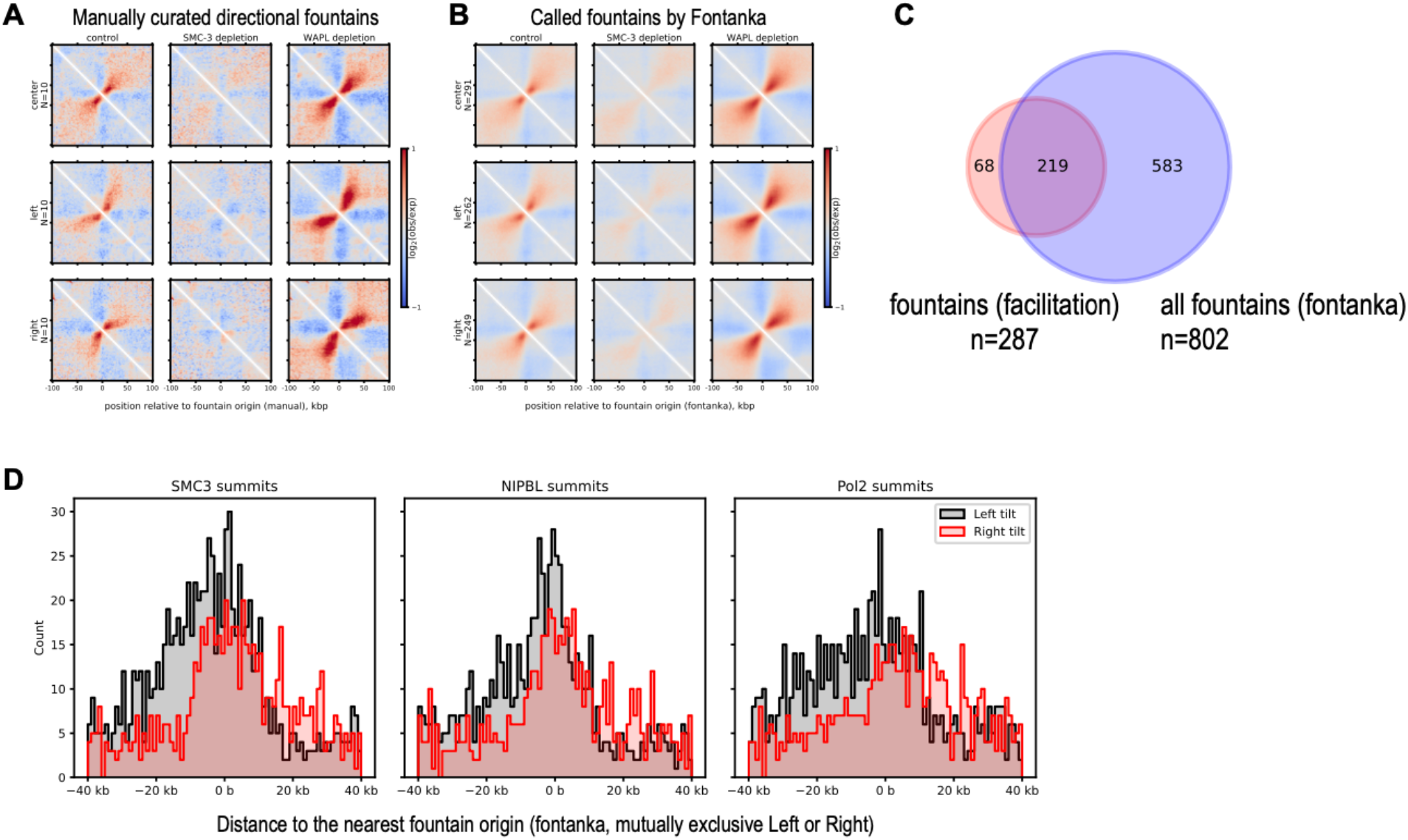
Identifying direction specific fountains using the fontanka tool. A) By manually scanning the genome, fountains are annotated left or right directed based on the orientation. B) Manually selected fountains are used as masks for identifying fountains following the published method (see Methods) (Galitsyna et al., 2023). The identified fountains are plotted for each group. C) Overlap between fountains identified by insulation scores in Supplemental Fig S1 and by fontanka. D) ChIP-seq summit distribution relative to the fontanka identified left and right tilted fountain origins.

**Supplemental Figure S11.**
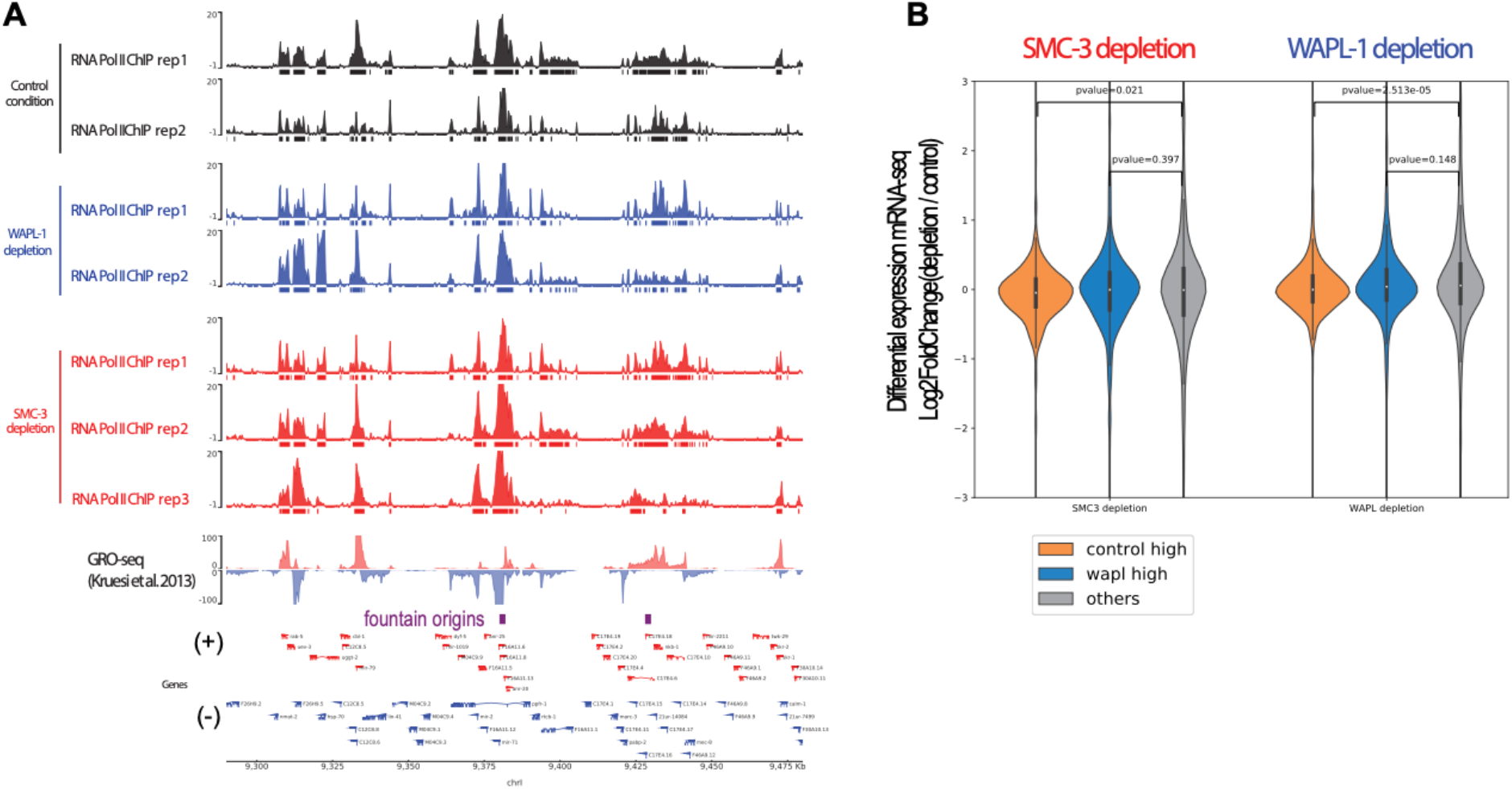
Replicates of RNA Pol II ChIP-seq data and mRNA-seq upon 4-hour depletion. A) RNA Pol II ChIP-seq tracks for the control, WAPl-1, and SMC-3 depletion conditions at an example region in the region. B) Differential expression values (log2fc from DESeq2) is plotted for three categories of genes upon 4-hour SMC-3 and WAPL-1 depletion. No-tag strain was used as control. Mann-Whitney test was used to generate p-values.

